# ARABIDOPSIS Bsister and SEEDSTICK MADS-box transcription factors modulate maternal nutrient flow for seed development in Arabidopsis

**DOI:** 10.1101/2025.07.21.665905

**Authors:** Camilla Banfi, Nicola Babolin, Chiara Astori, Chiara Mizzotti, Rosario Vega-Léon, Giulia Leo, Ueli Grossniklaus, Matthew R. Tucker, Fabrizio Araniti, Riccardo Aiese Cigliano, Walter Sanseverino, Ignacio Ezquer, Jose M. Muino, Kerstin Kauffman, Maurizio Di Marzo, Lucia Colombo

## Abstract

Successful seed development in angiosperms depends on the coordinated transport and allocation of sugars from maternal tissues to the developing embryo and endosperm. In *Arabidopsis thaliana*, ovules function as carbohydrate sink organs, accumulating starch in both gametophytic and sporophytic domains prior to fertilization. This stored starch is later mobilized to support early embryogenesis. Despite extensive knowledge of starch metabolism in photosynthetic tissues, the regulatory mechanisms governing sugar transport in reproductive organs remain poorly understood. Recent studies have identified fertilization-dependent changes in nutrient flow, including callose-mediated modulation of symplastic transport at the phloem unloading site. However, the molecular players orchestrating these transitions are largely unknown. Here, we show that the MIKC MADS domain transcription factors ABS/TT16 and STK play critical roles in regulating maternal nutrient flow during ovule maturation and seed development. We dissect their functional redundancy using omics and genetic approaches, underscoring the importance of different ovule tissues in coordinating sugar transport pathways for post-fertilization development. Our findings reveal a previously underappreciated layer of genetic control over nutrient allocation in reproductive tissues and provide new insights into the metabolic reprogramming required for successful seed formation.

## Introduction

Angiosperm ovules are complex structures in which the haploid female gametophyte (embryo sac) is surrounded by diploid sporophytic tissues, comprising the ovule integuments, nucellus, chalaza and the funiculus^1^. Arabidopsis has bitegmic ovules; each integument is initially composed of two layers named, from inside to out, inner integument 1 (ii1), inner integument 2 (ii2), outer integument 1 (oi1), and outer integument 2 (oi2)^2^. Before fertilization the ii1, termed endothelium, divides to form the fifth integument layer, called ii1’, which is located between the ii1 and ii2^1–3^. At ovule maturity, the integuments completely envelop the seven-celled female gametophyte containing the female gametes, the egg cell (EC) and the central cell (CC). In mature ovules, the residual nucellus and the chalaza are located between the female gametophyte and the funiculus, which connects the ovule to the placenta in the pistil. Double fertilization of the EC and the CC marks the conversion of ovules into seeds, after which the two fertilization products, the embryo and the endosperm, grow enclosed by the maternal seed coat^4^.

Arabidopsis ovules behave as sink organs, accumulating starch in gametophytic and sporophytic tissues^5–7^. During early stages of seed development, the starch contained in the embryo sac is metabolised and used as a nutritional resource to sustain early seed development^5^. Concurrently, starch accumulates in the seed chalaza and in the persistent nucellus^5,6^. Starch is a complex branched glucose polymer that serves as major storage carbohydrate in plants. Its formation relies on the translocation of hexose phosphate from the cytosol into amyloplasts. In ovules and seeds, starch is synthesized in the amyloplasts after long-distance sugar transport via the phloem. Sugars are transported from the placental tissue to the chalazal seed coat, where the vasculature ends (phloem end gate) and nutrient unloading takes place^8–12^. After unloading has occurred, the nutrient transport occurs either symplastically or apoplastically, eventually reaching the developing embryo and endosperm^10^. Recently, a fertilization-dependent mechanism regulating nutrient transport was identified in the seed, which depends on callose deposition/degradation at the phloem end gate^12^. Callose starts to be deposited in this region just before fertilization, and its removal allows symplastic nutrient transport to take place^12^. However, it has remained unclear how the sugar transport in the ovule and seed is regulated, and if it plays a role in the fertilization process and early stages of seed development.

*ARABIDOPSIS Bsister/TRANSPARENT TESTA 16* (*ABS/TT16*) and *SEEDSTICK* (*STK*) are sporophytically expressed MADS-box genes involved in several aspects of ovule and seed development. *ABS* regulates the differentiation and pigmentation of the endothelium, as well as nucellus degeneration^3,6,7,13–17^. In the *abs* mutant, the endothelium exhibits abnormal cell morphology, and after fertilization, fails to accumulate proanthocyanidins (PAs) that give the typical pigmentation to the Arabidopsis seeds^16^. In contrast to *abs*, the *stk* mutant is characterized by ectopic accumulation of PAs in the seed coat^18^. Before fertilization, *STK* controls ovule identity redundantly with *SHATTERPROOF 1* (*SHP1*) and *SHP2*^19^. *STK* is also required for seed coat differentiation, seed size and seed abscission^18–24^. In *abs stk* double mutants, ovules are characterized by an increased accumulation of starch, lack of endothelium differentiation and division, and severe fertility defects^7^.

We have integrated RNA sequencing (RNA-seq), Chromatin Immunoprecipitation Sequencing (ChIP-seq), and metabolomic profile, to unveil *ABS* and *STK* downstream pathways. We dissect *abs stk* phenotype, highlighting a compromised development of the embryo during the early stages of seed development, and we link this phenotype to the disrupted sugar metabolism and transport. Our results demonstrate that *ABS* and *STK* have pivotal importance in modulating the multistep maternal nutrient flow in developing ovules and seeds.

## Results

### Simultaneous loss of *ABS* and *STK* function causes ovule and seed defects

The *abs stk* double mutant, rather than the single mutants, significantly impacts ovule and seed development, causing partial sterility^3,7^. The *abs stk* ovules are characterized by the lack of ii1 periclinal division and differentiation, resulting in four-layered integuments (missing the ii1’ layer), a smaller embryo sac, and an increased ovule starch content^3,7^ (Fig. 1a). At the end of wild type (wt) ovule development, starch accumulates within the embryo sac, at the boundary between the nucellus and the chalaza, between the chalaza and the funiculus, and in the micropylar integuments (Fig. 1a). A similar pattern is observed in *abs* and *stk* single mutant ovules (Fig. 1a). By contrast, the *abs stk* ovules display higher starch accumulation in all these domains, especially inside the embryo sac and around the chalaza, consistent with impaired sugar metabolism (Fig. 1a). The starch enclosed inside the embryo sac gets degraded after fertilization in wt seeds, but persists in most *abs stk* seeds^7^. At 2 Days After Pollination (DAP), *abs stk* seeds are smaller in comparison to wt and *stk* and *abs* single mutants; moreover, they exhibit a pronounced nucellus (Fig. 1b). By counting and measuring oi2 cells, we found that the number of cells in *abs stk* is similar to that in *abs* mutants, whereas in terms of cell length, the *abs stk* double mutant resembles the *stk* single mutant (Fig. 1c). Most of the *abs stk* seeds eventually abort at different stages, resulting in only 11% of viable seeds at maturity^7^.

**Fig. 1.**
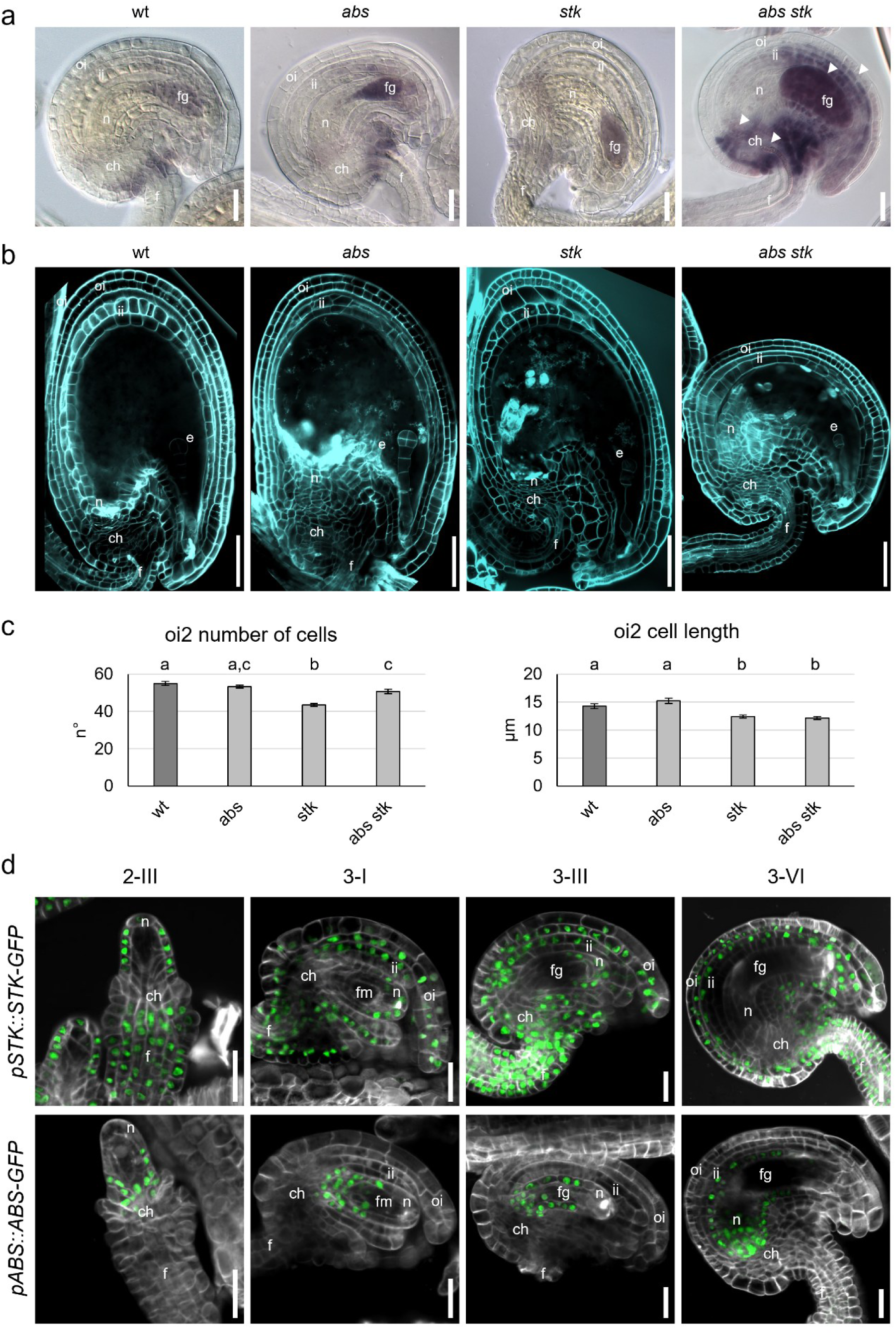
*abs stk* mutant is characterized by increased starch accumulation in ovules and a defective seed coat. **a** Microscopy images of wt, *abs*, *stk* and *abs stk* mature ovules stained with Lugol’s solution that marks starch accumulation (purple). The arrowheads highlight the overaccumulation of starch in the *abs stk* mutant. **b** Confocal images of wt, *abs*, *stk* and *abs stk* seeds at 2 DAP. Seeds were stained with SR2200 (cyan). **c** Graphs showing the number of cells composing the oi2 and the length of the cells composing the oi2 of wt (n=8), *abs* (n=8), *stk* (n=8) and *abs stk* (n=12) seeds at 2 DAP. Error bars represent the standard error mean. In each graph, different letters above the bars represent significant differences (*p* < 0.05) as determined by one-way ANOVA followed by Tuckey’s HSD, whereas the same letters indicate absence of a statistical difference between the respective genotypes. Two independent measurements showed similar results. **d** Expression of *pSTK::STK-GFP* and *pABS::ABS-GFP* during ovule development. Cell walls are stained with SR2200 (white). ch: chalaza, e: embryo, f: funiculus, fg: female gametophyte, fm: functional megaspore, ii: inner integument, n: nucellus, oi: outer integument. Scale bars 20 µm (**a, d**), 50 µm (**b**). Experiments were repeated on four independent individuals with similar results, and representative images are shown in **a, b, d**.

To determine ABS and STK protein localization during ovule development, we used functional *pSTK::STK-GFP* and *pABS::ABS-GFP* marker lines^3,18^. Both fusion proteins are expressed only in the sporophytic tissues of the ovule (Fig. 1d) and seed^3^. In young ovules at stage 2-III, we detected STK-GFP in the upper nucellus, in the basal part of the chalaza, and the funiculus. Later, at stages 3-I/3-III, the GFP signal was detected in the outer integuments and the outermost layer of the inner integuments (oi1, oi2, ii2), in the upper nucellus, until its degeneration at stage 3-VI, and in the funiculus. ABS-GFP was localized in the lower part of the nucellus (ovules stage 2-III, 3-I, 3-III), in which it persisted until the mature ovule stage, and in the two innermost layers of the inner integuments (ii1 and ii1’) at stages 3-VI (Fig. 1d). These data reveal striking complementarity of the expression patterns for ABS and STK, suggesting the two proteins do not need to be expressed in the same cell types to jointly regulate ovule development.

### Transcriptome analysis in *abs stk* mutants highlights deregulation of genes related to sugar metabolism

To identify the molecular networks controlled by *STK* and *ABS* in ovules before fertilization, we performed RNA sequencing (RNA-seq) of mRNA from isolated pistils of wt, *abs*, *stk*, and *abs stk* mutants (Supplementary Fig. 1a). Differential expression analysis identified differentially expressed genes (DEGs) between the wt and the three mutant backgrounds (Fig. 2a, Supplementary Fig. 1b). The double and single mutants were also compared to identify genes with unique or additive expression changes (Fig. 2a, Supplementary Fig. 1b). KEGG (Kyoto Encyclopedia of Genes and Genomes) pathway enrichment analysis revealed common enriched pathways in *abs*, *stk* and *abs stk* to wt comparisons. Among the common enriched categories there were metabolic pathways (ath01100), biosynthesis of secondary metabolites (ath01110), carbon metabolism (ath01200) and starch and sucrose metabolism (ath00500). These data are consistent with the previously described roles of *ABS* and *STK* in the flavonoid pathway ^14,18^, and in cell wall synthesis and modification^21,22,24^. Furthermore, the enrichment of the starch and sugar metabolism term (ath00500) in the *abs stk* to wt comparison is consistent with the excess of starch in double mutant ovules (Fig. 2b). Considering the gene ontology results and the *abs stk* phenotypes, we focused our attention on genes involved in starch metabolism and sugar transport. In the *abs stk* to wt comparison, the most upregulated gene in starch biosynthesis was *GRANULE-BOUND STARCH SYNTHESE 1* (*GBSS1*), which encodes an enzyme required in the initial phase of starch biosynthesis (Fig. 2c)^25^. On the other hand, *β-AMYLASE 4* (*BAM4*) was the most downregulated gene involved in starch degradation^26^. Among the genes coding for the sugar transporters, *GLUCOSE 6-PHOSPHATE/PHOSPHATE TRANSLOCATOR-2 (GPT2*) was highly upregulated in *abs stk*, whereas *SUCROSE-PROTON SYMPORTER 5* (*SUC5*), encoding a sucrose transporter, was the strongest downregulated gene (Fig. 2c)^27–29^. Starch metabolism was reported to be influenced by trehalose, a disaccharide associated with physiological effects on growth and carbon allocation in plants^30^. Most of the genes involved in trehalose synthesis, belonging to the *TREHALOSE-6-PHOSPHATE SYNTHASE* (*TPS*) and *TREHALOSE-6-PHOSPHATE PHOSPHATASE (TPP)* gene families, were downregulated in the *abs stk* double mutant compared to wt (Fig. 2c). However, the most differentially expressed genes were *TPPI*, which was upregulated in the double mutant, and *TREHALASE 1* (*TRE1*), the only gene involved in trehalose degradation, which was downregulated^31^. All the 35 genes with altered expression in *abs stk* flowers compared to wt, are usually expressed in mature unfertilized wt ovules according to the transcriptomic data produced by Zhang and collaborators^32^. To verify the expression changes in the double mutant ovules, we have analysed *TRE1* and *BAM4* expression by *in situ* hybridization (Supplementary Fig. 2). Consistent with the transcriptomic data, we detected lower levels of both transcripts in *abs stk* compared to wt, especially in ovule sporophytic tissues.

**Fig. 2.**
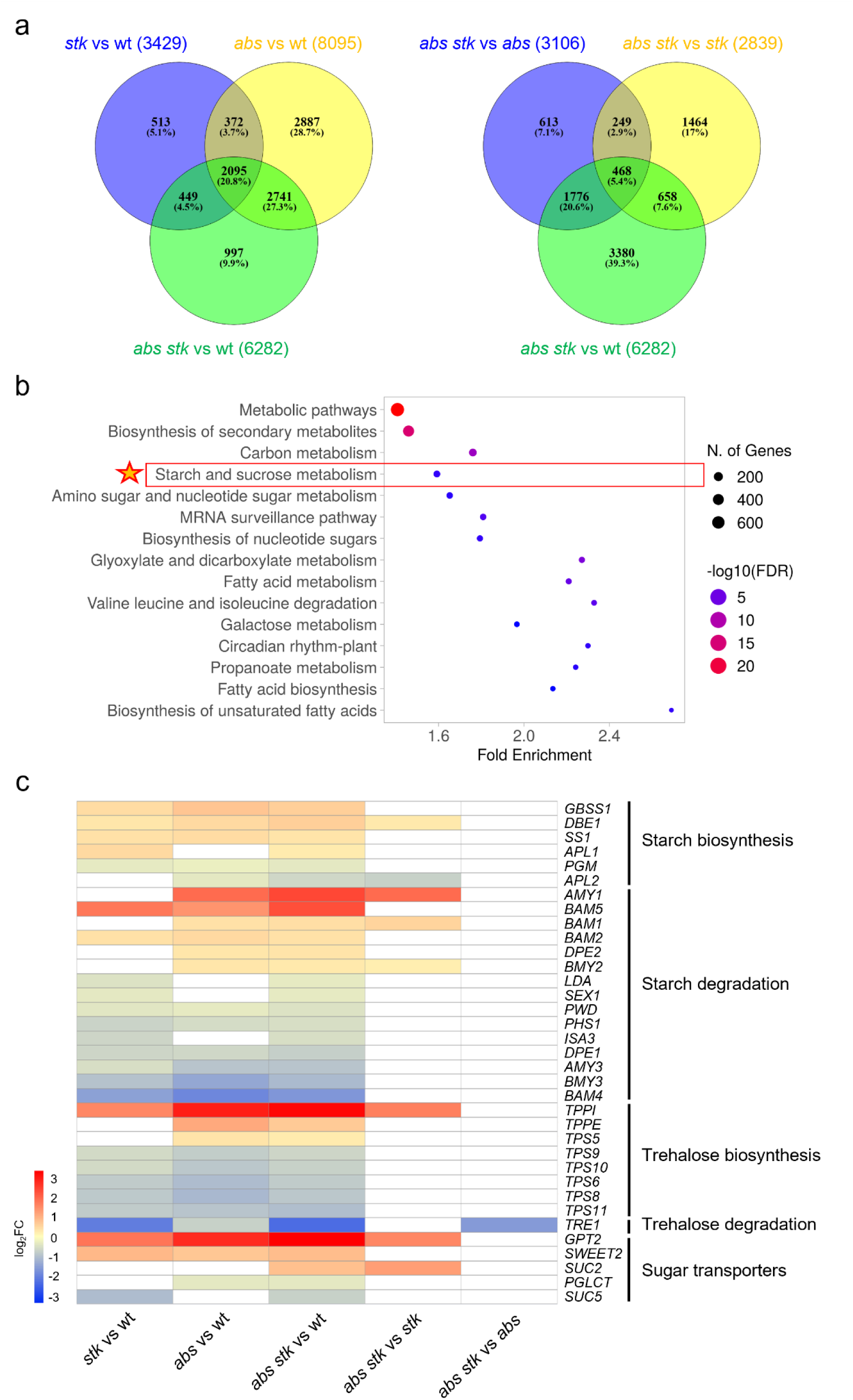
Starch and sucrose metabolic genes are deregulated in the *abs stk* double mutant before fertilization. **a** Venn diagrams of the DEGs in mutant to wt comparisons and in the double to single mutant comparisons. FDR ≤ 0.05 was considered to identify the DEGs. n=4. **b** Dot plot of KEGG pathway enrichment analysis showing the 15 most enriched pathways in the *abs stk* to wt comparison. The starch and sucrose metabolism pathway is highlighted. The enriched KEGG pathways are ordered by the number of genes in each enriched category. FDR < 0.05 was considered for significance. **c** Heat map showing the DEGs involved in starch and trehalose metabolism, and sugar transport, in the mutant to wt comparisons and in the double to single mutant comparisons. The genes shown in this table were filtered for the DEGs in *abs stk* to wt comparison. Inside each category, the genes are ordered for the log_2_FC in the *abs stk* to wt comparison from the highest (red) to the lowest (blue).

The analysis of the DEGs related to sugar and starch metabolism suggests that there is no additive effect in the double mutant with respect to the single mutants. Of the 35 DEGs between *abs stk* and wt, 26 were differentially expressed in *stk* and 29 in *abs* compared to wt, suggesting that *ABS* and *STK* act redundantly on many genes in these pathways (Fig. 2c). In conclusion, these findings indicate that *ABS* and *STK* modulate the expression of genes involved in starch and trehalose metabolism in ovules.

### *STK* and *ABS* loss-of-function mutations impact sugar content

To evaluate if the differences in gene expression in *abs stk* mutants correlate with metabolic alterations, we performed a Gas Chromatography/Mass Spectrometry (GC/MS)-driven metabolomic analysis on *abs, stk,* and *abs stk* mutant flowers before anthesis. The untargeted metabolomic analysis allowed the extraction of 323 compounds, of which 135 were annotated (Supplementary Fig. 3a and b). Among them, soluble amino acids and sugars were some of the classes characterized by the highest VIP (Variable Importance in Projection) score values (Supplementary Fig. 3c). Univariate analysis revealed that 122 out of 135 metabolites significantly differed between the wt and at least one of the three mutants (Supplementary Fig. 4). Considering the *abs stk* starch phenotype and the deregulation of genes involved in starch metabolism in the double and single mutants compared to the wt found in the RNA-seq analysis, we focused attention on soluble sugars. Four of these, including sucrose, glucose 6-phosphate, fructose 6-phosphate, and trehalose, were more abundant in the *abs stk* double mutant compared to the wt and at least one of the single mutants (Fig. 3a). Sucrose is the major transport form for photo-assimilated carbon and the major source of energy for non-photosynthetic tissues like ovules and seeds^33^. We found significantly higher sucrose accumulation in the *abs* and *abs stk* mutants compared to wt, whereas *stk* showed lower sucrose accumulation (Fig. 3a). The hydrolysis of sucrose produces two important monosaccharides, glucose and fructose, that, upon phosphorylation, provide carbon for cell wall and starch biosynthesis^34^. The accumulation patterns of glucose 6-phosphate and fructose 6-phosphate were similar to that of sucrose, with lower levels in the *stk* mutant and higher levels in *abs* and *abs stk* compared to the wt (Fig. 3a). The level of trehalose was additive in the double mutant with respect to the two single mutants, as the highest level of this compound was present in the *abs stk* double mutant, followed by *abs* and *stk* single mutants and the wt (Fig. 3a). Thus, ABS appears to be a key factor limiting accumulation of soluble sugars in young flowers, whereas STK tends to promote their accumulation. *abs* is epistatic to *stk* in terms of sucrose, glucose 6-phosphate and fructose 6-phosphate accumulation. In contrast, the two genes act together to limit trehalose accumulation, since levels are additively increased in *abs stk* mutants.

**Fig. 3.**
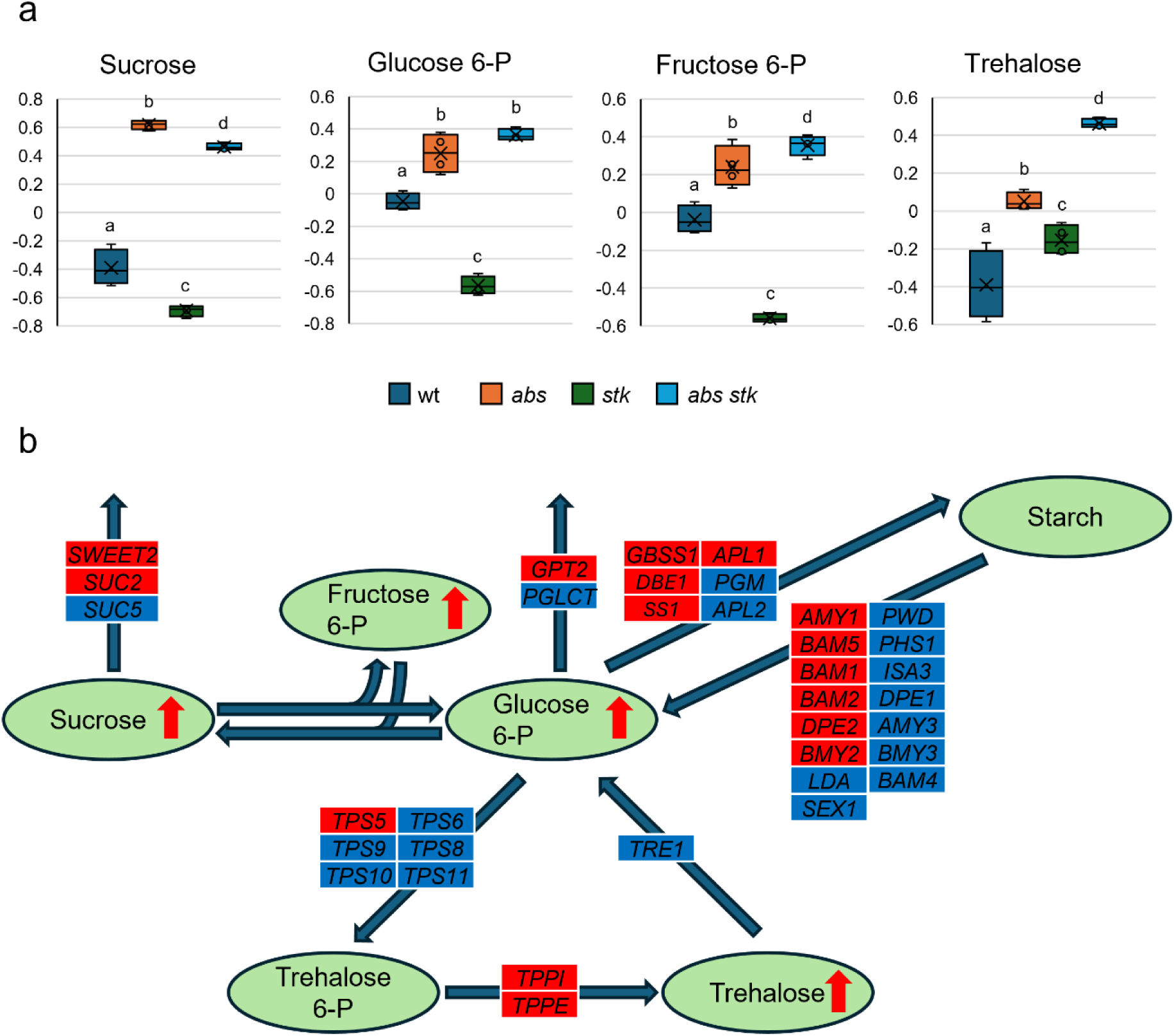
Sugar accumulation is perturbed in the *abs stk* mutant. **a** Graphs showing the quantification of sucrose, glucose 6-P, fructose 6-P and trehalose in wt, *abs*, *stk* and *abs stk* flowers by GC-MS. In each graph, different letters above the bars represent significant differences (*p* < 0.05) as determined by univariate one-way ANOVA using LSD test as post-hoc, whereas the same letters indicate absence of a statistical difference between the respective genotypes. n=4. **B** Schematic representation of alterations in the starch and sucrose metabolic pathway in the *abs stk* to wt comparison integrating the transcriptomic and metabolomic data. The coloured rectangles represent the DEGs in *abs stk* to wt comparison involved in each step of the pathway; downregulated genes are indicated in blue, whereas upregulated genes are indicated in red. The arrows in the circles, representing the metabolites, indicate the altered quantity of the metabolite in the *abs stk* to wt comparison. Red arrows indicate sugars with a significantly higher quantity in *abs stk* mutant.

By integrating the results of transcriptomic and metabolomic analyses, we generated a graphical representation of the key metabolic nodes related to the aforementioned sugars, highlighting the metabolic alterations in *abs stk* with respect to wt (Fig. 3b). The high starch content in the double mutant may not be solely attributed to elevated levels of the glucose 6-phosphate precursor, that was present at the same level in the *abs* mutant (Fig. 3a), but also from the upregulation of several genes involved in starch biosynthesis and the downregulation of genes responsible for its degradation (Fig. 3b). Similarly, increased trehalose, which was consistent with upregulation of *TPP* genes, mediating the dephosphorylation of trehalose-6-phosphate, coupled with a marked downregulation of *TRE1*, may contribute to regulate starch accumulation^30^ (Fig. 3b).

The integration of omics data suggests that the altered level of starch in *abs stk* double mutant ovules possibly derives from the deregulation of multiple categories of ovule-expressed genes, that cause an overall sugar balance modification.

### STK directly regulates sugar metabolism-related genes

MADS-box transcription factors regulate transcription by binding consensus sequences, CC(A/T)_6_GG, termed CArG box, in the regulatory region of their target genes^35^. To gain deeper insights into the possible involvement of the MADS-box TF STK in the control of sugar metabolism genes, we performed Chromatin immunoprecipitation sequencing (ChIP-seq) experiment using inflorescences before anthesis of *pSTK::STK-GFP* plants (Fig. 1d). We obtained 6799 significant peaks mapped to the reference genome with a total of 5485 genes that were directly bound by STK. Overall, 93% of the peaks were in the promoter region of the 5486 putative STK target genes (Supplementary Fig. 5a). Functional enrichment analysis (Gene Ontology, GO) of target genes revealed that the most significant categories of the molecular functions were transcription factor activity, sequence-specific DNA binding (GO:0003700) and DNA binding (GO:0003677). Interesting data were obtained from the analysis of the GO terms of the biological processes. In this case, the enriched categories were seed coat (GO:0010214), fruit development (GO:0010154), gynoecium development (GO:0048467), and flavonoid biosynthetic process (GO:0009813), consistent with previously reported roles of *STK*^18,22,36–38^. Other significant categories included carbohydrate metabolism, such as response to sucrose (GO:0009744), carbohydrate transport (GO:0008643), carbohydrate biosynthetic process (GO:0016051), and trehalose biosynthetic process (GO:0005992), suggesting a possible involvement of this TF in directly regulating sugar metabolism. Finally, one of the most enriched categories in the analysis of KEGG pathways was starch and sucrose metabolism (ath00500) (Supplementary Fig. 5b). In particular, STK was found to bind the genomic sequences of one starch biosynthetic gene, eight genes involved in starch degradation, twelve genes involved in trehalose metabolism, and eleven genes encoding sugar transporters. By comparing this gene list with the RNA-seq data, we found that most of the STK targets were downregulated in *abs stk* compared to the wt (10/12), while only two were upregulated. Together, these identify STK as a direct regulator of several genes involved in sugar metabolism, mostly acting as a transcriptional activator.

### Modulation of starch metabolism partially rescues *abs stk* fertility defects

To evaluate whether the *abs stk* defects could be due to the lacking capability to degrade starch in the female gametophyte, required for early stages of embryo development, we expressed *BAM5*, encoding a starch-degrading enzyme, under the control of the CC-specific *pDD22* promoter^39^. *BAM5* is not usually expressed in the CC in wt^40^. By lugol staining of ovules, we confirmed that the introgression of the *pDD22::BAM5* construct decreases starch accumulation in the female gametophyte of the wt and *abs stk* mutants (Fig. 4a). Surprisingly, we observed a rescue of the ii1 periclinal division in the *abs stk pDD22::BAM5* line, resulting in five-layered ovules at maturity (Fig. 4b). To evaluate the percentage of fertilized ovules, we crossed the wt and the mutant female plants, with the *pWOX9::WOX9-GFP* reporter line as pollen donor^41^. At 3 DAP, we observed a higher percentage of embryos expressing WOX9-GFP in *abs stk pDD22::BAM5* (71%) compared to *abs stk* plants (55%) (Supplementary Fig. 6a and b). Furthermore, analysis of developing seeds in the siliques of three independent *abs stk pDD22::BAM5* lines showed an increase in seed viability up to 64% compared to 14% in *abs stk* double mutant (Fig. 4c, Supplementary Fig. 6c).

**Fig. 4.**
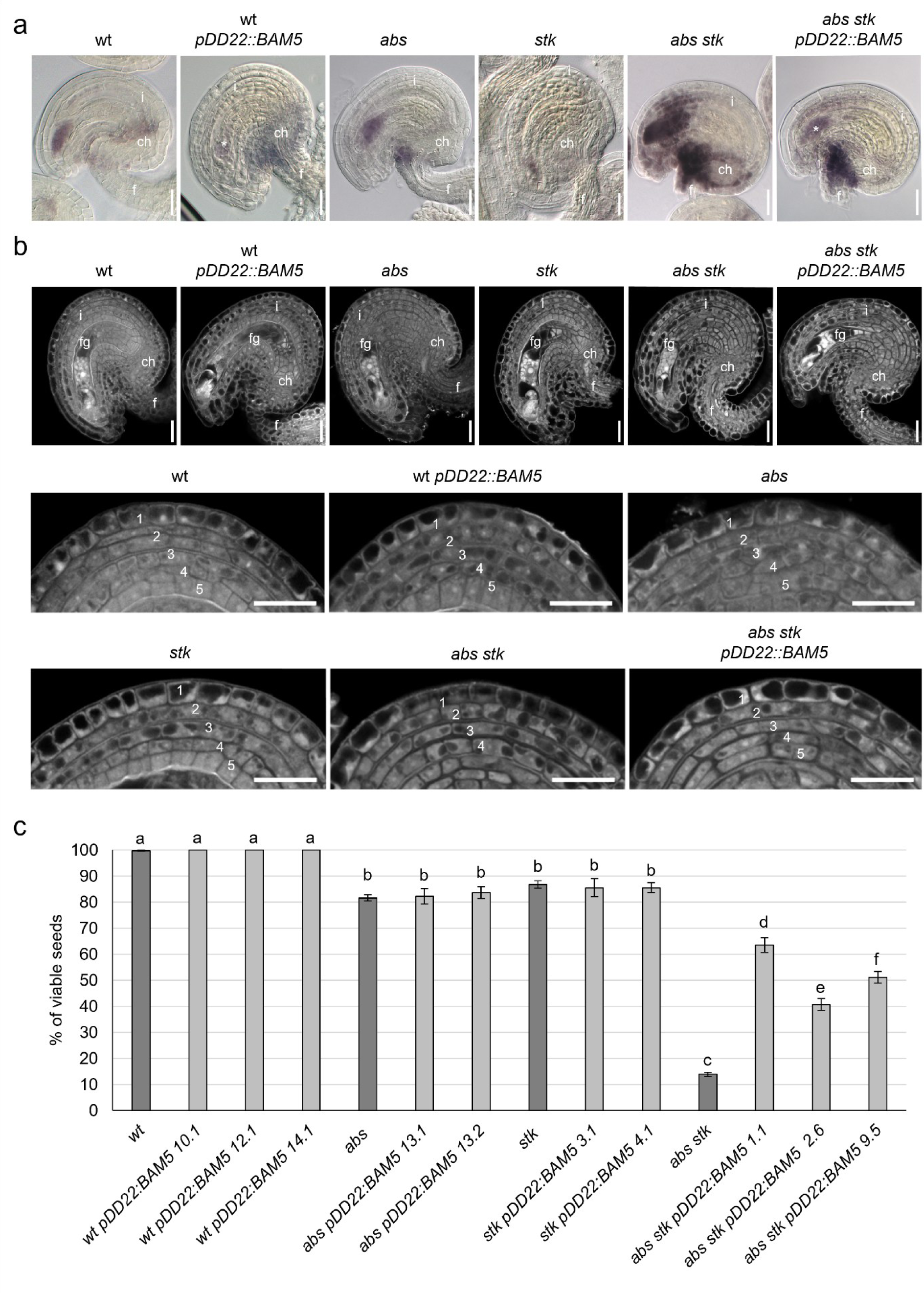
Starch degradation in *abs stk* central cell partially rescues seed formation and integument defects. **A** Microscopy images of wt, wt *pDD22::BAM5*, *abs*, *stk*, *abs stk,* and *abs stk pDD22::BAM5* mature ovules stained with Lugol’s solution that marks starch accumulation (purple). The asterisks indicate the reduced starch accumulation caused by *pDD22::BAM5* expression. **B** Confocal images of wt, wt *pDD22::BAM5*, *abs*, *stk*, *abs stk,* and *abs stk pDD22::BAM5* mature ovules, and magnification of the ovule integuments showing the number of integument layers. Ovules were stained by Feulgen staining. **C** Graph showing the percentage of viable seeds in siliques of wt (n=19), wt *pDD22::BAM5* 10.1 (n=8), wt *pDD22::BAM5* 12.1 (n=8), wt *pDD22::BAM5* 14.1 (n=8), *abs* (n=20), *abs pDD22::BAM5* 13.1 (n=5), *abs pDD22::BAM5* 13.2 (n=5), *stk* (n=24), *stk pDD22::BAM5* 3.1 (n=7), *stk pDD22::BAM5* 4.1 (n=6), *abs stk* (n=26), *abs stk pDD22::BAM5* 1.1 (n=15), *abs stk pDD22::BAM5* 2.6 (n=14), and *abs stk pDD22::BAM5* 9.5 (n=15). Error bars represent the standard error mean. Different letters above the bars represent significant differences (*p* < 0.05) as determined by one-way ANOVA followed by Tuckey’s HSD, whereas the same letters indicate absence of statistical difference between the respective genotypes. Measurements were repeated two times. Ch: chalaza, f: funiculus, fg: female gametophyte, i: integument. Scales bars 20 µm. Experiments were repeated on four independent individuals with similar results, and representative images are shown in **a, b**.

Since at 3 DAP the *abs stk* phenotype was not fully rescued by *pDD22::BAM5* expression, we monitored zygote and embryo formation in seeds at 1 and 2 DAP by confocal laser scanning microscopy. In wt seeds at 1 DAP, the zygote had an elongated shape, before it divided by asymmetrical cell division, resulting in the embryo-proper reaching the 2/4-cell stage at 2 DAP (Fig. 5a)^42,43^. At 1 DAP, the percentage of seeds bearing zygotes was not significantly different among all the analysed genotypes, indicating that the *abs stk* is not defective in EC fertilization (Fig. 5b). However, at 2 DAP, the development of *abs stk* embryos was delayed; we observed a higher percentage of seeds still containing zygotes (35%) in comparison to all other analysed genotypes (Fig. 5b). Remarkably, the *pDD22::BAM5* construct rescued early embryo development in *abs stk*, whereby the percentage of developing zygotes and embryos at 2 DAP is similar to the wt (Fig. 5b). In conclusion, in the *abs stk* mutant, zygotes likely degenerate or stop developing just after fertilization, consistent with the WOX9-GFP embryonic marker being expressed in just ∼50% of the *abs stk* seeds at 3 DAP (Supplementary Fig. 6a and b).

**Fig. 5.**
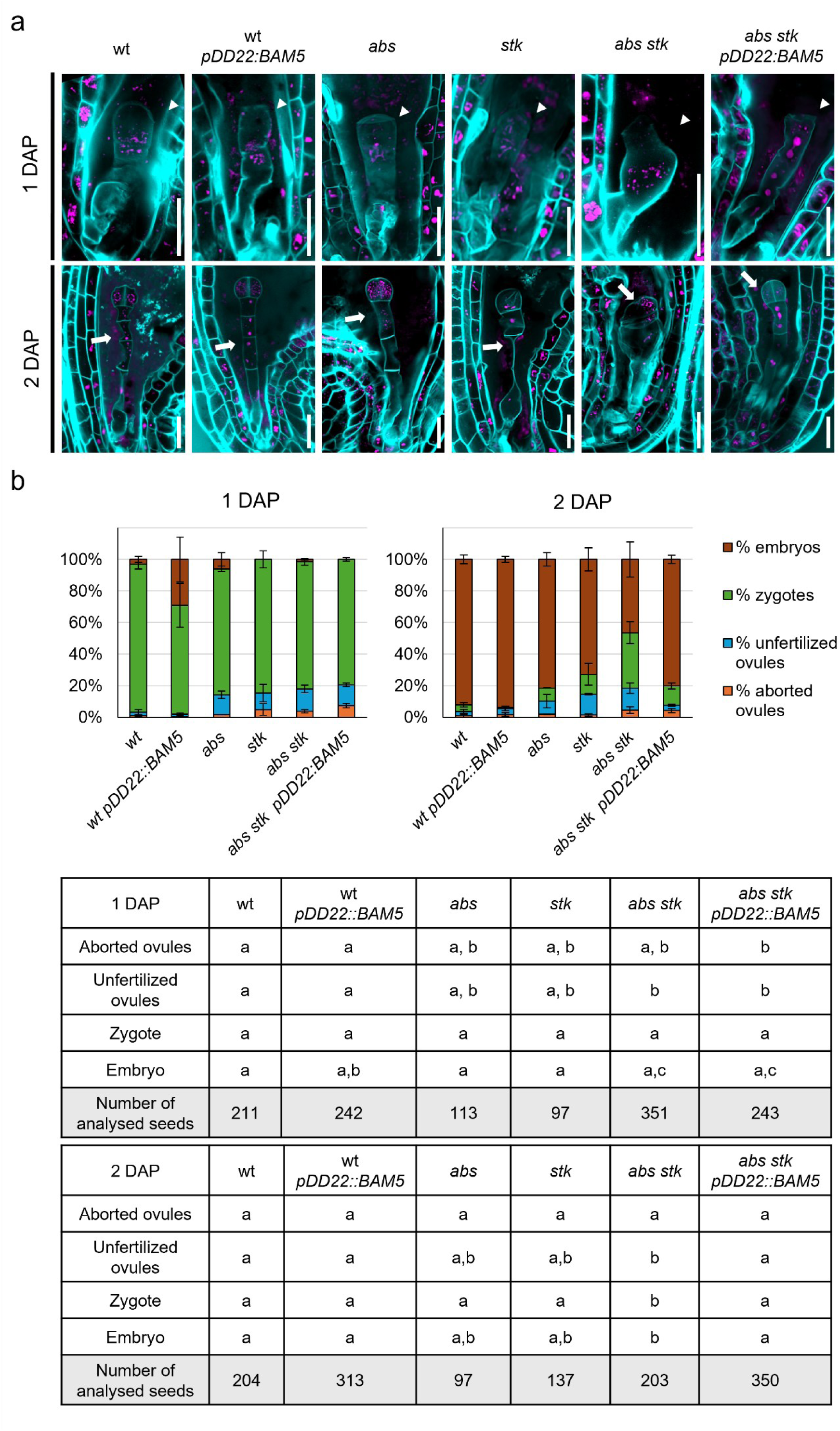
Degradation of starch in *abs stk* central cell rescues embryo development. **A** Confocal images of wt, wt *pDD22::BAM5*, *abs*, *stk*, *abs stk,* and *abs stk pDD22::BAM5* seeds at 1 DAP and 2 DAP showing the presence of zygotes and embryos. Arrowheads point the zygotes, while arrows point the embryos. Seeds were stained with SR2200 (cyan), and propidium iodide (purple). Scale bars 20 µm. **b** Graphs showing the percentage of aborted ovules, unfertilized ovules, seeds bearing zygotes, and seeds bearing embryos observed at 1 and 2 DAP by confocal analysis. Error bars represent the standard error mean. In each row of the tables, different letters above represent significant differences (*p* < 0.05) as determined by one-way ANOVA followed by Tuckey’s HSD, whereas the same letters indicate absence of statistical difference between the respective genotypes. The total number of analysed ovules/seeds for each genotype and stage is indicated in the table. At least two independent measurements showed similar results. Experiments were repeated on four independent individuals with similar results, and representative images are shown in **a**.

Collectively, these data demonstrate that the simultaneous absence of *ABS* and *STK* in ovule sporophytic tissues impacts the early stages of embryo development. This seems to be caused by the altered availability or regulation of sugars required to sustain the growth of the early embryo, as embryo development could be partially restored by degrading the starch in the central cell and fertilization-derived endosperm.

### Callose turnover is disrupted in the phloem unloading domain of *abs stk* seeds after fertilization

Although the *abs stk* fertility and seed set defects were partially rescued by degrading the starch inside the female gametophyte, in the *abs stk pDD22::BAM5* rescue lines, seed set was still significantly lower in comparison to the wt (Fig. 4c). One possible explanation for this difference is that starch breakdown in the CC increases free sugars to sustain the early stages of zygote growth and division, but additional sporophytic defects might still be present in the *abs stk pDD22::BAM5* lines. Since sugars typically move from the sporophytic tissue to the embryo sac, we analysed the nutrient flow during early seed development by using the 5(6)-Carboxyfluorescein diacetate (CFDA), a fluorescent molecule transported through the phloem vasculature and subsequently into the ovule through symplastic connections. In all genetic backgrounds, the fluorescent tracer is visible in the funiculus and the chalaza, but not in the integuments of unfertilized ovules at 1 Day After Emasculation (1 DAE), indicating the absence of symplastic connectivity (Fig. 6a). However, at 3 DAP, we detected the CFDA tracer in the outer layers of the seed coat in wt, wt *pDD22::BAM5*, *stk* and *abs* single mutants, as symplastic movement is restored after fertilization (Fig. 6a)^8^. Conversely, in the *abs stk* double mutant and in *abs stk pDD22::BAM5* seeds, CFDA migration was still blocked in the chalazal seed coat, consistent with compromised nutrient flow during early seed development (Fig. 6a). Recently, it has been demonstrated that the nutrient flow into seeds after fertilization is regulated by callose degradation at the phloem end gate^12^. We thus investigated callose accumulation in ovules and seeds by aniline blue staining. In wt, wt *pDD22::BAM5,* and *stk* and *abs* single mutants, we observed callose deposited at the phloem end gate at 1 DAE, whereas at 3 DAP callose was reduced (Fig. 6b). Although at 1 DAE no major differences were observed in *abs stk* compared to the wt and the single mutants (Fig. 6b), in *abs stk* and in *abs stk pDD22::BAM5* seeds at 3 DAP, the callose deposition was still present at the phloem pole, likely blocking symplastic flow (Fig. 6b).

**Fig. 6.**
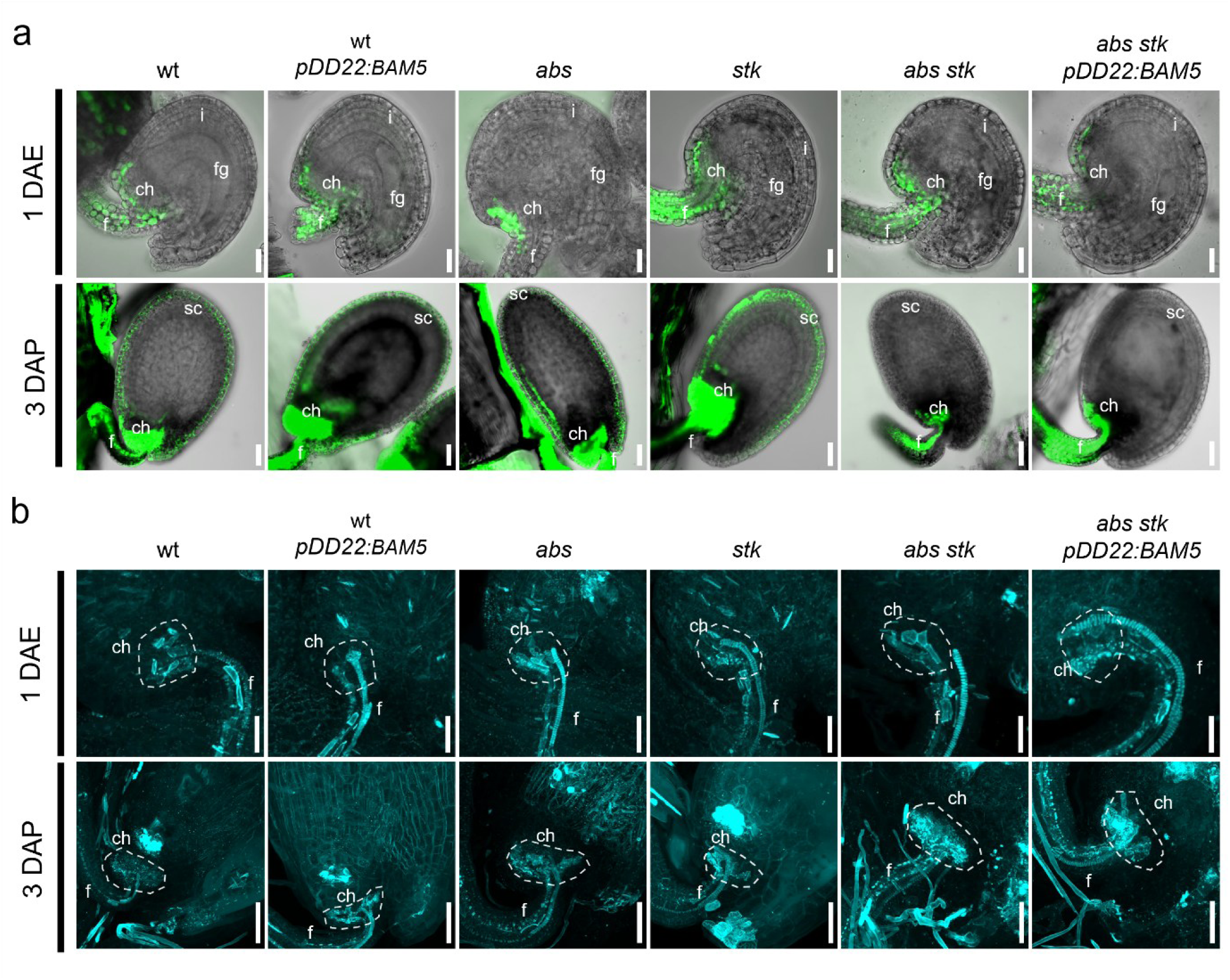
*Abs stk* early seeds have defects in nutrient flow and callose deposition. **A** Confocal images of ovules at 1 DAE and seeds at 3 DAP showing CFDA symplastic tracer accumulation (green) in wt, wt *pDD22::BAM5*, *abs*, *stk*, *abs stk,* and *abs stk pDD22::BAM5* mature ovules at 1 DAE and seeds at 3 DAP. The GFP channel is merged with the brightfield channel to highlight the ovules and seeds profile. **B** 3D reconstruction of Z-stacks confocal images of mature ovules at 1 DAE and developing seeds at 3 DAP stained with aniline blue (cyan), highlighting callose deposition at the phloem end gate, indicated by dashed line. ch: chalaza, f: funiculus, fg: female gametophyte, i: integuments, sc: seed coat. Scale bar 20 µm (ovules 1 DAE), 50 µm (seeds 3 DAP). Experiments were repeated on three independent individuals with similar results, and representative images are shown in **a, b**.

These results suggest that in the *abs stk* double mutant, sap flow from the sporophytic tissues to the embryo sac is impeded at the early stages of embryo development.

### STK and ABS control callose gate opening after fertilization to ensure proper seed development

The absence of callose removal after fertilization is consistent with blocked movement of CFDA and nutrient flow in the *abs stk* double mutant seeds, but not in the single mutants. This unique phenotype prompted us to examine genes whose expression was deregulated in the *abs stk* double mutants compared to the single mutants and wt. To this aim, we performed an RNA-seq experiment on siliques of all genotypes at 3 DAP, including the *abs stk pDD22::BAM5* and wt *pDD22::BAM5* lines (Fig. 7a, Supplementary Fig.7a and b). Recently, Liu and collaborators revised the number of genes predicted to degrade callose to a total of forty-two^12^. On the other hand, only twelve genes for callose synthesis have been reported in Arabidopsis^44^. Among the DEGs in the *abs stk* double mutant compared to the wt, we identified eleven genes encoding callose-degrading enzymes, but no genes involved in callose biosynthesis (Fig. 7b). Notably, ten out of eleven genes involved in callose degradation were downregulated in the *abs stk* mutant compared to wt (Fig. 7b), of which two (AT2G01630, AT1G32860) are usually expressed in the chalazal seed coat domain according to snRNA-seq ^45^ and thus may be directly involved in the absence of callose removal in *abs stk* seeds. Genes such as *beta-1,3-glucanase 3* (*BG3*) and *BG1*, which are essential for callose degradation, had the lowest expression in the *abs stk* double mutant compared to both wt and the single mutants (Fig. 7b), but they are not expressed in the seed chalazal region in wt condition. Some of these *BGs* were downregulated in the *abs* mutant, but not in the *stk* single mutant compared to wt (Fig. 7b), suggesting concerted activity of multiple hydrolytic genes may be required to remove the callose plug.

**Fig. 7.**
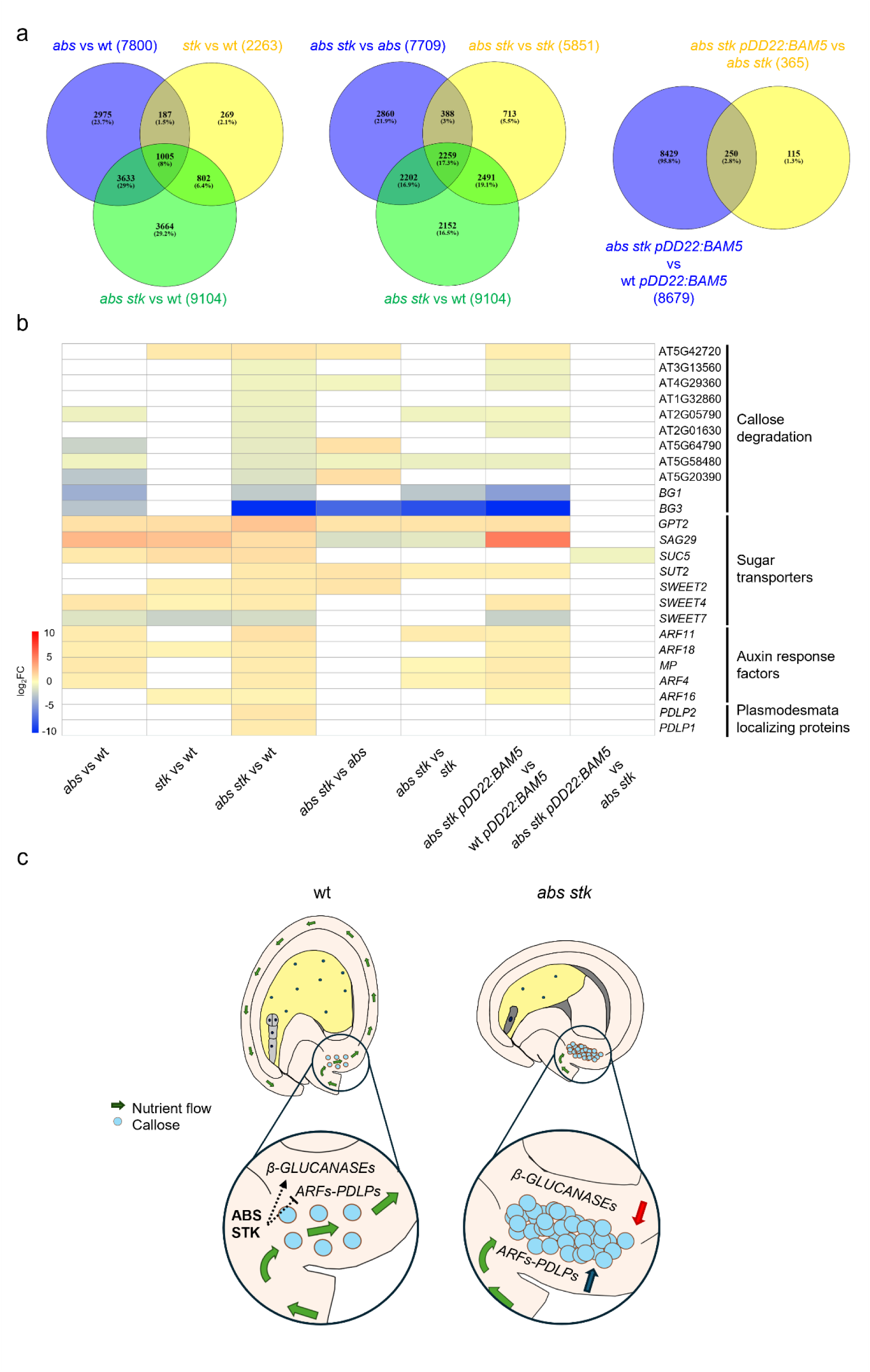
Genes involved in callose turnover, sugar transport, and auxin response are deregulated after fertilization by *ABS* and *STK* mutations. **A** Venn diagrams of the DEGs in the mutants to wt comparisons, in the double to single mutant comparisons and in the *abs stk pDD22:BAM5* comparisons. FDR < 0.05 was considered to identify the DEGs. n=4. **b** Heat map showing the DEGs involved in callose metabolism, sugar transport and auxin response in the mutant to wt comparisons, in the double to single mutants comparisons and in the in the *abs stk pDD22:BAM5* comparisons. The genes shown in this table were filtered for the DEGs in *abs stk* to wt comparison. Inside each category, the genes are ordered for the log_2_FC in *abs stk* to wt comparison. Colors represent the log_2_FC values, from the highest (red) to the lowest (blue). **c** Putative model representing ABS and STK activity in regulating callose deposition and nutrient flow in the chalazal seed coat.

Given that nutrient flow is impaired in the double mutant, we further examined the expression of genes encoding sugar transporters, including the families of *SUGARS WILL EVENTUALLY BE EXPORTED TRANSPORTER (SWEET)* and *SUC*. The *SWEET* family encodes seventeen transporters involved in sugar movement, and we identified three *SWEET*s that were upregulated in the *abs stk* mutant in comparison to wt, while only *SWEET7* was downregulated. Also, among the nine *SUC* genes, two were upregulated in the double mutant compared to the wt (Fig. 7b). However, none of these differentially expressed genes is usually expressed in the seed coat chalazal domain^45^, consistent with our hypothesis that the symplastic flow in *abs stk* seed chalaza is affected, rather than the apoplastic transport. Recent reports indicated a key role of auxin in modulating plasmodesmata aperture through the interplay of the *AUXIN RESPONSE FACTORS* (*ARFs*) and the *PLASMODESMATA-LOCATED PROTEIN* (*PDLP*) in roots^46,47^. Therefore, we considered whether a similar mechanism might be involved in callose deposition in the chalazal seed coat. Out of twenty-three *ARFs*, *ARF4*, *MONOPTEROS* (*MP*/*ARF5*), *ARF11*, *ARF16*, and *ARF18* were upregulated in the *abs stk* mutant compared to the wt (Fig. 7b). Among them, *MP* is also upregulated in *abs* single mutant compared to wt, while *ARF16* is upregulated in *stk* vs wt, suggesting some specificity of ABS and STK in the regulation of *ARF* genes (Fig. 7b). Regarding *PDLPs*, we found *PDLP1* and *PDLP2* were upregulated in *abs stk* with respect to the wt, but, notably, not in the single mutant vs wt (Fig. 7b) comparisons. Interestingly, only the expression of *MP, ARF18* and *PDLP1* overlaps in the chalazal seed coat in wt condition^45^ (Supplementary Fig. 8a and b), where callose deposition occurs, suggesting a potential role for these two proteins in modulating callose deposition to regulate nutrient flow into the developing seed, or a non-cell autonomous regulation mediated by other factors. Lastly, most of the DEGs identified in the comparison of *abs stk pDD22::BAM5* to wt *pDD22:BAM* showed a similar trend when comparing *abs stk* to wt, suggesting that the rescue mediated by the *pDD22::BAM5* construct is not related to callose turnover, sugar transport and the ARF-PDLP pathway, but likely only to starch degradation (Fig. 7b).

*STK* expression domain, but not *ABS*, partially overlap with the chalazal seed coat domain in seeds at 3 DAP^45^. To assess how many of these callose and solute transport-related genes might be direct targets of STK, we performed ChIP-seq on developing fertilized siliques of *pSTK::STK-GFP*. This revealed 8231 significant peaks, primarily located in the promoter regions of the genes (Supplementary Fig. 9), with a total of 6287 genes directly bound by STK. We identified 13 callose turnover-associated genes targeted by STK after fertilization, however, only one (AT5G20390), which is not expressed in the seed chalazal domain^45^, was downregulated in *abs stk* compared to wt in the RNA-seq data. This suggests that ABS might be the primary regulator of *BGs* expression, or intermediate factors regulated by *STK* and *ABS* act upstream of the callose hydrolytic machinery. Consistent with this hypothesis, only 4 *ARFs,* including *ARF4, MP, ARF11* and *ARF16,* and one *PDLP* (*PDLP2*), were found to be either direct targets of STK and upregulated in *abs stk* mutant compared to wt. This suggests that after fertilization, STK appears to be part of a mechanism that regulates the expression of nutrient flux-related genes. Taken together, our findings suggests that the callose degradation at the phloem end gate, which regulates nutrient flux in the chalazal seed coat, is controlled by ABS and STK, potentially through a non-cell autonomous complex network including genes encoding callose-degrading enzymes, together with *ARF*-*PDLP* modules (Fig. 7c).

## Discussion

Maternal sporophytic tissues play a crucial role during seed development, embedding embryo and endosperm, and thus the next generation. The crosstalk between the ovule’s sporophytic tissues and the embryo sac is fundamental for the development of ovules and seeds, and the ovule integuments are a critical part of this communication^48–50^. In *abs stk* ovules, the inner integument fails to differentiate correctly, and endothelium identity is lost^7^. Moreover, *abs stk* mutants show high seed sterility, suggesting the two phenotypes might be linked. Although the inner integument is needed for the correct development of a mature female gametophyte^51^, genetic ablation of the endothelium in ovules and seeds still results in the formation of viable embryo and endosperm^16,51^. In *abs stk* ovules, no apparent defects are observed in female gametophyte formation, suggesting that the two layers of the inner integument are sufficient for gametogenesis. Furthermore, our data presented here suggest that the developmental defects observed during embryogenesis in the double mutant are likely independent of the lack of the endothelium differentiation. Hence, the basis for the post-fertilization defects in *abs stk*, and the pathways involved, may have been overlooked in previous studies. Other than the lack of endothelium, the double mutant presents a range of phenotypes. For example, *abs stk* seeds possess smaller seed coat cells and impaired seed coat expansion compared to wt (Fig. 1). This phenotype can be attributed to the *stk* mutation, which has previously been associated with smaller outer integument cells in the seed coat^23^. Nevertheless, the *stk* mutation alone is not responsible for the fertility defects that characterize the *abs stk* double mutant, as the percentage of viable seeds in the double mutant is significantly lower than in the *stk* single mutant (Fig. 4). In addition, young seeds of *abs stk* show a lack of nucellus degeneration, but this phenotype is also observed in *abs* mutants that still show high levels of seed fertility^52^. Again, these data point towards other ABS- and STK-dependent ovule pathways being compromised that contribute to early seed development.

Although the *abs stk* ovules harbor a mature gametophyte, the lack of *ABS* and *STK* function in the sporophytic tissues impacts the development of embryo and endosperm^7^ (Fig. 1, 4). These phenotypes are of sporophytic maternal origin^53^, as the double heterozygotes *abs/ABS stk/STK* plants, or even the *abs/ABS stk/stk* or *abs/abs stk/STK* plants, do not present the same fertility defects^7^. In barley and wheat, the Bsister MADS-box gene *MADS31* is sporophytically expressed in the ovule nucellus, which differentiates to become the primary solute transfer tissue in the grain. *mads31* mutants exhibit defects in nucellus differentiation, partial defects in gametophyte development, and show reduced seed set^54^. Even if the *abs* mutation alone does not affect gametophyte development and seed set in Arabidopsis, the role of Bsister genes in controlling communication between the sporophytic and gametophytic generations may be conserved. A previous study showed that at 3 DAP, only half of the *abs stk* seeds contain a developing endosperm; however, the percentage of developing embryos had not been investigated^7^. Here, by analysing the presence of zygotes, we demonstrated that the EC is correctly fertilized in *abs stk* seeds (Fig. 5). However, embryo development is delayed and arrests at later stages (Supplementary Fig. 6). It has been shown that mutations in sucrose transporter encoding genes (*SWEET4/11/12*/*15* and *SUC5*), which are expressed in the persistent nucellus and seed coat, cause a delay in embryo development including a reduction of the size of the embryo and the whole seed (Baud et al., 2005; Chen et al., 2015, Lu et al., 2021). Nonetheless, the *sweet11/12/15* and *suc5* mutants form a normal embryo. Most of the sugar transporter-encoding genes are upregulated or expressed normally in *abs stk* mutant compared to wt, suggesting that other factors are likely involved in embryo arrest.

Many genes encoding proteins of the starch and sucrose metabolic pathways are deregulated in the *abs stk* double mutant, possibly contributing to the starch over-accumulation observed in double mutant ovules (Fig. 1, 2). Previous studies showed that *STK* is involved in the modulation of key sugar metabolism genes during seed coat development ^21,22^, but no information is available for *ABS*. Our data demonstrate that STK is a direct transcriptional regulator of genes involved in starch breakdown, trehalose and sugar transport. Members of the glucan water dikinase (*GWD*) and amylase (*AMY* and *BAM*) gene families play a crucial role in catalyzing the hydrolysis of this polyglycan into maltose units^25^. *GWD1* and *BAM4*, direct targets of STK, are downregulated in the *abs stk* double mutant and in the single mutants compared to wt (Fig. 2). However, only the *abs stk* double mutant accumulated more starch in ovules, and this phenotype is likely caused by a general imbalance of the sugar metabolic pathways. We also report higher levels of trehalose in the *abs stk* mutant in comparison with the wt and the single mutants (Fig. 3). Higher trehalose levels may inhibit starch degradation and induce its over accumulation^55^. The downregulation of *TRE1*, a direct STK target, in the *abs stk* double mutant possibly contributes to higher trehalose accumulation and, in turn, increases the quantity of starch (Fig. 2).

ABS and STK interact with SEP3 in yeast, suggesting they may form a complex^56^. However, the localization of STK and ABS proteins during all stages of ovule development indicates that they are located in different ovule domains. *STK* is mainly expressed in the outer integuments and the funiculus, whereas *ABS* is detected in the nucellus and the two inner integuments (Fig. 1). RNA-seq and ChIP-seq data indicate that both genes are involved in the regulation of the sugar metabolic pathway pre-fertilization. This regulation should occur, at least partially, in a non-cell autonomous manner, as *ABS* and *STK* are not detected in the embryo sac where the largest starch deposit is present in the *abs stk* mutant. Some of the STK direct targets are also regulated by ABS, as they are differentially expressed in the *abs* single mutant in comparison to the wt. Nevertheless, other metabolic genes are differentially expressed exclusively in one of the two single mutants compared to wt, suggesting that these two TFs could regulate genes from the same pathway in a redundant manner, but in different domains of ovules and seeds. Consistent with this hypothesis, in seeds, nutrient transporters belonging to the UMAMITs, for amino acids, and SWEETs, for sucrose, specifically mark distinct seed compartments^10,11,57^. From these domains, they transport nutrients from the sporophytic tissues to the fertilization products and since multiple transporters exist, the mutation of single genes does not compromise seed viability^10^.

Our results suggest that the fine-tuned pre- and post-fertilization sugar balance has a major impact on early seed development. Moreover, this balance relies on the communication between the sporophyte and gametophyte. By reducing the starch in the embryo sac of *abs stk* seeds we could rescue the early stages of embryo development (Fig. 4 and 5). In wt seeds, as soon as double fertilization takes place, the starch accumulated in the CC is broken down, likely mobilizing nutrients to sustain the developing embryo, and potentially allowing for further influx of sugars. In *abs stk*, starch over-accumulates in the embryo sac. This suggests that sugar transport pathways are functional prior to fertilization, but turnover appears to be compromised. Similarly, starch is maintained unused after fertilization^7^, and fertility is only restored through exogenous expression of *BAM*. Starch degradation may therefore increase the level of free sugars to support embryo growth just after fertilization. Sugars are transported to the ovules and seeds by the phloem tissues, which end in the chalazal region at the phloem end gate. Here they are symplastically unloaded just after fertilization and transported to the other seed domains^8^. The phloem unloading of CFDA, a marker for symplastic transport, occurs in wt and single mutant seeds but is blocked in the *abs stk* double mutant (Fig. 6). The absence of nutrient transport from the phloem end gate to the embryo sac likely causes starvation that compromises the early stages of zygote development in *abs stk* seeds. Recently, a strong relationship has been demonstrated between sugar signalling and cell cycle progression^58^; this supports our hypothesis that the altered sugar accumulation and metabolism in *abs stk* seeds may contribute to early stages of embryo division and growth. Considering this, our data also demonstrates how enhancing the sugar availability in the embryo sac induces division in the endothelial cell layer (Fig. 4).

Recently, the nutrient supply to the seed has been related to callose gate removal at the phloem end gate in response to successful fertilization. In the fertilization-defective *generative cell-specific 1* (*gcs1*) mutant, nutrient flow is not triggered due to increased callose accumulation, preventing the waste of nutrients^12^. Callose over-accumulates at the phloem end gate of *abs stk* seeds, blocking nutrient transport, and the starch mobilization in the embryo sac is not sufficient to induce callose removal (Fig. 6). Similarly, *AtBG_ppap* loss-of-function mutation, disrupting a putative plasmodesmata-associated β-1,3-glucanase (a callose-degrading enzyme), causes increased callose accumulation in the chalazal seed domain and smaller seeds. However, in the *Atbg_ppap* mutants, the transport of nutrients is not completely compromised, and seed viability is not affected, indicating that other redundant genes may control callose gate opening at the phloem end gate^12^. Our results indicate that many genes involved in callose degradation are downregulated in the *abs stk* double mutant after fertilization, possibly contributing to impaired callose removal (Fig. 7). Notably, one of the most downregulated genes in the *abs stk* double mutant is *BG1*, which has already been linked to the deficiency in callose degradation observed in the *gcs1* mutant^12^. However, according to snRNA-seq data^45^, *BG1* is not expressed in the seed coat chalazal domain, suggesting it could act in a non-cell autonomous way, or most likely other β-1,3-glucanases may regulate callose gate opening. Furthermore, it has been reported that auxin plays a role in modulating plasmodesmata aperture through the interplay between the *ARFs* and *PDLPs*^46,47,59^. Plants overexpressing *PDLP5* are characterized by increased callose levels and suppressed plasmodesmal trafficking^47^. In *abs stk* double mutant, the upregulation of *ARF* and *PDLP* genes, normally expressed in the chalazal domain of the seed, might contribute to the impermeability of plasmodesmata by callose deposition, thus blocking nutrient flux to the seed (Fig. 7). Although our results suggest that *ABS* and *STK* directly or indirectly regulate callose deposition via *BGs* and *ARF-PDLP* modules, further analyses are needed to identify the signal(s) that induces callose degradation and opening of the phloem end gate, and the reason why this signal is not perceived in *abs stk* seeds.

Our findings demonstrate the importance of STK and ABS, two MADS-box transcription factors, in controlling ovule development and the flux of solutes between the maternal sporophyte and the next generation. The two proteins appear to act in concert from different ovule tissues to support sugar accumulation and metabolism, which is required for embryo growth, and to subsequently remove callose barriers adjoining the nucellus and seed coat that are required for post-fertilization seed development. Thus, ABS and STK provide a critical regulatory link between maternal nutrient supply and seed developmental processes.

## Methods

### Plant material and growth conditions

All plants used in this study were of the Columbia-0 (Col-0) accession. Seeds were sown either directly on soil or on Murashige and Skoog (MS) medium^60^ and then moved to soil. Before sowing in germination medium, seeds were surface-sterilized with 70% ethanol for 2 minutes (min), 1% bleach for 5 min, and then washed three times with sterile water. After 10 days, the seedlings were moved to soil and grown in a growth chamber under long-day conditions (22°C, 16 h light/8 h dark). The *abs* (*tt16-6*) and *stk* (*stk-2*) mutants were already described^19,61^. Mutants were screened using PCR. The *pSTK::STK-GFP* marker line used in this work was previously generated^18^. The *pABS::ABS-GFP* was kindly provided by E. Magnani and previously described^3^, and the *pWOX9::WOX9-GFP* was previously described^41^.

### Plasmid construction and Arabidopsis transformation

The expression vector pK2GW7 was modified by Doulix (Venice, Italy) to substitute the *p35S* followed by the Gateway cassette with the synthetically generated *pDD22* promoter upstream of the CDS of *BAM5*. For the promoter, 935 bp before ATG were considered, as described previously^39^. The resulting expression vectors were transformed into *Agrobacterium tumefaciens* by electroporation, and Arabidopsis plants were transformed using the floral dip method^62^. Seeds of transformed plants were selected on kanamycin (100 mg L^−1^) on MS medium. The presence of the construct was assessed by PCR. Three independent T1 lines were selected, and the T2 plants were further analysed.

### Fertility analyses

The analysis of fertilization efficiency was performed by opening mature siliques of wt, *abs*, *stk*, *abs stk*, wt *pDD22::BAM5*, *abs pDD22::BAM5, stk pDD22::BAM5,* and *abs stk pDD22::BAM5,* counting the number of viable seeds per silique using a Leica MZ6 stereomicroscope. Images were acquired and processed using Axiovision (version 4.1) software. To confirm the data obtained by the fertility analysis, we used a marker line containing the *pWOX9::WOX9-GFP* construct by crossing these plants (used as pollen donors) with wt *pDD22::BAM5*, *abs pDD22::BAM5, stk pDD22::BAM5, abs stk pDD22::BAM5,* and *abs stk*. The derived siliques were collected at 3 DAP, dissected in water on a glass slide to obtain seeds, and finally observed using a Nikon A1 confocal laser scanning microscope to count the number of seeds expressing the GFP signal. GFP was excited with a 488 nm laser, and emission was detected between 520 and 540 nm. Confocal images were analysed using Fiji software^63^. The statistical differences were calculated using one-way ANOVA with post-hoc Tukey’s HSD test (*p* < 0.05).

### Feulgen staining of ovules

For the morphological characterization of ovules, pistils were isolated from flowers before anthesis^64^. The reagents and protocol used to perform Feulgen staining were described previously^65^. The samples were excited with a laser at 488 nm and emission was detected between 515 and 600 nm. Images were captured using a Nikon A1 confocal laser scanning microscope. Confocal images were analysed using Fiji software^63^.

### Lugol staining of ovules

To visualize starch accumulation, pistils were isolated from flowers before anthesis^64^. Starch staining was performed with Lugol’s solution as previously described^7^. We then dissected the pistils in water to isolate the ovules. The signal was observed using a Zeiss Axiophot D1 microscope equipped with DIC optics. Images were recorded with an Axiocam MRc5 camera (Zeiss) using the Axiovision program (version 4.1).

### Expression analysis by *in situ* hybridization

*In situ* hybridization and probe synthesis had been performed as described previously^66^. Arabidopsis inflorescences before anthesis were sampled, fixed, and embedded in paraffin. Plant tissue sections (8 μm) were probed with antisense digoxigenin (DIG)-labelled probes. The signal was observed using a Zeiss Axiophot D1 microscope equipped with DIC optics. Images were recorded with an Axiocam MRc5 camera (Zeiss) using the Axiovision program (version 4.1).

### RNA-seq material and data analysis

Total RNA was extracted from a pool of isolated pistils from stage 9 to stage 12^64^ of wt, *abs*, *stk*, and *abs stk*, in four biological replicates, using the Macherey Nagel “Nucleospin RNA Plant” kit following the manufacturer’s instructions (pre-fertilization samples). For post-fertilisation samples, RNA was extracted in the same way from a pool of 3 DAP siliques of wt, *abs*, *stk*, *abs stk*, wt *pDD22:BAM5* and *abs stk pDD22:BAM5*, in four biological replicates. mRNA-seq with stranded oligo-dT specific library preparation and sequencing of 20 million of PE150 reads was performed by Biomarker Technologies (BMK) with Illumina NovaSeq X platform. The obtained raw reads were trimmed with Trimmomatic 0.39^67^, with the following parameters: ILLUMINACLIP:1:30:6, LEADING: 0, TRAILING: 20, AVGQUAL: 20, MINLEN: 35. STAR 2.7.10b^68^ was used for mapping, with --alignIntronMax 10000 on the TAIR10 genome. Counts were obtained using featureCounts 2.1.1^69^, using Araport11 annotation, with the following parameters: -p, - -countReadPairs. HTSFilter^70^ was used to filter the counts. Principal component analysis (PCA) was performed to identify clustering between the four biological replicates of each sample. DESeq2^71^ was exploited to obtain the DEGs. Only DEGs with an FDR ≤ 0.05 were considered as statistically significant. DEGs analysis was performed on R studio, version 4.4.1. Venn Diagrams were generated with the online tool Venny 2.1.0. The functional enrichment analysis of RNA-seq (KEGG) was performed using the online tools ShinyGO 0.81 with an FDR cut-off value of 0.05 to select the enriched categories.

### ChIP-seq material and data analysis

ChIP-seq was performed as described previously^72,73^ with the following modifications. Approximately 1 g of inflorescences with flowers until stage 12 (pre-fertilization) and siliques from stage 14 to stage 16 (post-fertilization) of *stk* plants complemented with *pSTK::STK-GFP*^18^, in three biological replicates, were crosslinked with 1% (v/v) formaldehyde under vacuum for 2x 15 min, with mixing in between. The crosslinking reaction was stopped by adding glycine to 0.25 M concentration. The fixed tissue was ground in liquid nitrogen followed by nuclei isolation and chromatin extraction. The chromatin was sonicated using the S220 (Covaris) sonicator with the following settings: Peak Power = 140.0; Cycles/Burst = 200; Duty Factor = 5.0; Duration = 590 sec; temp = 6°C. Samples were centrifugated and a portion of the sonicated chromatin (supernatant) was set aside as input control. 5 µl of anti-GFP antibody (Abcam, ab290) was added to the sonicated chromatin and mixed at 4°C for 1 h. A portion of µMACS Protein A MicroBeads (Miltenyi Biotec) was blocked with 0.5% BSA in IP buffer mixing at 4 °C for 6 h and added to the mix. The immunoprecipitated chromatin was captured mixing at 4 °C for 1 h. The samples were passed through a magnetic µColumn (Miltenyi Biotec) fixed on a µMACS Separator (Miltenyi Biotec) and washed three times with each IP buffer, high-salt buffer, LiCl buffer, and TE buffer. The immunoprecipitated chromatin was eluted with hot elution buffer (1% SDS, 0.1M NaHCO_3_). To reverse the crosslinking, input and eluate samples were incubated with 250 mM NaCl in a thermomixer at 65 °C for 16 h and 1000 rpm, followed by incubation with 0.615 mg/ml proteinase K at 65 °C for 1 h and 1000 rpm. The de-crosslinked DNA was ethanol-precipitated and purified using the DNA Clean & Concentrator kit (Zymo Research). The ChIP-seq library was prepared with the ThruPLEX DNA-seq Kit (Takara Bio) following the manufacturer’s instructions. The DNA libraries were size selected for 100-700 bp fragments with the AMPure XP beads (Beckman Coulter). The size-selected libraries were assessed on a 4200 TapeStation (Agilent) and the concentration measured using a Qubit 4 instrument (Invitrogen). The libraries were sequenced using Illumina high-throughput sequencing device.

The quality of the raw sequencing reads was assessed with *FastQC*. Potential adapter sequences were removed from the sequencing reads using *Trimmomatic*^67^ and the quality of trimmed reads was reassessed. Trimmed reads were mapped to the Arabidopsis TAIR10 reference genome using *Bowtie2*^74^. Only reads with MAPQ score > 40 were kept. Peak calling and fragment pileup was performed using *Macs2*^75^ with parameters: *-p 0.05 -B –SPMR –mfold 2 20 -g 101274395*. For the STK-GFP ChIP-seq, the regions declared significant by *Macs2* were merged using *mergeBED* from the *bedtools* package^76^ with parameters: *-iobuf 5G -d 100* to obtain a common reference set to later test with *DESeq2*^71^. Finally, the R package ChIPseeker (annotatePeak) was used to annotate the peaks on the reference genome with a distance to TSS of −3000 and +3000 bp^77,78^. The functional enrichment analysis of ChIP-seq (GO and KEGG) was performed using the online tools DAVID with an FDR cut-off value of 0.05 to select the enriched categories.

### Extraction, identification, and quantification of primary metabolites in Arabidopsis pistils

The dissected pistils from flowers before fertilization (from stage 11 to 12^79^) were collected at 12.00 am and immediately frozen in liquid nitrogen to quench the metabolism. Freshly homogenized (50 mg) pistils were obtained for each biological replicate (4) and genotype (Col-0, *abs*, *stk*, and *abs stk*). Metabolite extraction and derivatization for untargeted metabolomics were carried out following the protocols described by Lisec et al. (2006)^80^. The derivatized extracts were injected into a 5MS capillary column (30 m x 0.25 mm x 0.25 µm) (with 10 m of pre-column) (Thermo Fisher Scientific, Waltham, MA, USA) using a single quadrupole GC/MS. Injector and source were set at 250 °C and 260 °C. 1 µl of the sample was injected in a splitless mode with a helium flow of 1 ml/min using the following programmed temperature: isothermal 5 min at 70 °C followed by a 5 °C/m ramp to 350 °C and a final 5 min heating at 330 °C. Mass spectra were recorded in electronic impact (EI) mode at 70 eV, scanning at 40-600 m/z range, scan time 0.2 second (s). The mass spectrometric solvent delay was set at 9 min. Pooled samples that served as quality control (QCs), n-alkane standards (C8-C40 all-even), and blanks were injected at scheduled intervals for instrumental performance, tentative identification, and monitoring of shifts in retention indices (RI). The RAW files obtained from the GC/MS analyses were analysed using MS-DIAL version 4.20 (http://prime.psc.riken.jp/compms/msdial/main.html)^81^. The centroided EI-MS data with at least 20 data points across the peaks were considered for further deconvolution using a sigma window value of 0.5. Spectral annotation was performed using an open-source EI-spectra reference assembled as described by Misra (2019). A retention index (RI) reference generated in-house with n-alkanes as described above was used for RI matching, and a cosine similarity of 70% was used as a cut-off for spectral matching with a reference database for compound ID assignment. For metabolite annotation, we followed the Metabolomics Standards Initiative (MSI) levels of the International Metabolomics Society: annotations were considered level 2 (putative annotation based on spectral library similarity) or level 3 (putatively characterized compound class based on spectral similarity to known compounds of a chemical class as suggested (Sumner et al., 2007). Spectral features found in blank runs were filtered out when present > 2 fold higher than in samples on average. For relative quantification purposes, when we encountered multiply silylated (n-TMS) features of well-annotated metabolites, we retained the major (higher abundant) compounds and left out other minor (low abundance) versions for consistent comparison across all samples. A matrix of annotated metabolites and their corresponding abundances across all samples were used for further processing, summary statistics, and data visualisation. The metabolomic experiments were carried out adopting a completely randomised design with five replications. Statistical analysis for the GC/MS dataset was performed using MetaboAnalyst version 4.0^82^. Briefly, relative abundance values from the MS-DIAL outputs were normalised using the QC and the internal standard ribitol as the reference standard, then were Log transformed and pareto scaled. Normalised data were successively statistically analysed through univariate (one-way ANOVA using LSD test as post-hoc, *p* < 0.05) and multivariate analysis [unsupervised principal component (PCA) analysis, and supervised PLS-DA analysis]. To avoid overfitting the PLS-DA model was validated using Q2 as a performance measure, the 10-fold cross validation and setting in the permutation test a permutation number of 20. Data were further represented through a heatmap, built on the top 60 compounds resulting from the ANOVA analysis, and hierarchically clustered using the Euclidean distance measurement and Ward as a clustering algorithm.

### Confocal microscopy of embryos

To analyse zygotes and embryos, siliques from hand-pollinated pistils were collected at 1 DAP and 2 DAP. Samples were treated with 0.2 M NaOH 1% (v/v) SDS solution for 3 h at 37°C, rinsed three times with distilled water, and finally moved in 2% (v/v) bleach solution at room temperature for 10 min. Siliques were rinsed five times with distilled water, then dissected using needles on a glass slide, and seeds were stained with SCRI Renaissance 2200 (SR2200) and Propidium Iodide (PI) solution (Merk). Images were acquired using a Nikon A1 confocal laser scanning microscope. SR2200 was excited with a 405 nm laser line, and emission was detected between 440 and 470 nm, whereas PI was excited with a 561 nm laser line and emission was detected between 660 and 730 nm. Confocal images were analysed using Fiji software^63^.

### Aniline blue staining of ovules and seeds

Emasculated pistils (1 DAE) and hand-pollinated siliques (3 DAP) were treated as reported previously^12^. Samples were collected in 1 M NaOH solution and left overnight at room temperature. Afterwards, the samples were incubated in aniline blue solution [0.1% (w/v) aniline blue and 0.1 M K_3_PO_4_] for at least 3 hours. Ovules/seeds were isolated with needles on a glass slide, and observed using a Nikon A1 confocal laser scanning microscope. The samples were excited with a laser at 445 nm and emission was detected between 492 and 545 nm. Z-stack images were acquired using a Nikon A1 confocal laser scanning microscope (Z-step 0.5 µm). 3D images were obtained using the 3D project tools on Fiji software^63^.

### CFDA tracer detection in ovules and seeds

Pistils (1 DAE) and hand-pollinated siliques (3 DAP) were cut with a sharp blade at the root of the silique pedicel and placed in PCR tubes containing 100 μl of 5(6)-Carboxyfluorescein diacetate (CFDA) solution (0.1 mg/ml, Merck) at 22℃ for 4 h^12^. The pistils and siliques were dissected with needles on a glass slide under a stereomicroscope to isolate ovules and seeds and mounted in water. Images were acquired using a Nikon A1 confocal laser scanning microscope. The CFDA was excited with a laser at 488 nm, and emission was detected between 520 and 540 nm. Confocal images were analysed using Fiji software^63^.

### Seed coat cells measurements

Siliques derived from hand-pollinated pistils (2 DAP) were treated as described in the “Confocal microscopy of embryos” paragraph. Seeds were stained with SCRI Renaissance 2200 (SR2200). Images of seeds were acquired using Nikon A1 laser scanning confocal microscope. SR2200 was excited with a 405 nm laser line, and emission was detected between 440 and 470 nm. Confocal images were analysed using Fiji software^63^, and cells of the oi2 layer were measured and counted, considering at least 8 seeds for each genetic background. Statistical differences between the different genotypes were analysed using one-way ANOVA with post-hoc Tukey’s HSD test (*p* < 0.05).

### Statistics and reproducibility

Apart from RNA-seq, ChIP-seq and metabolome profile, described in the Methods section, statistical analysis was performed using one-way ANOVA with post-hoc Tukey’s Honestly Significant Difference (HSD) test with the following webtools: https://astatsa.com/OneWay_Anova_with_TukeyHSD/ or https://www.statskingdom.com/180Anova1way.html. *p*-value < 0.05 was interpreted as statistically significant unless otherwise stated. Data are presented as mean ± Standard Error of Mean (SEM). Other details such as sample size and the level of significance are specified in the text, and figure legends. The experiments shown in the paper were repeated at least three times, unless otherwise stated.

## Data availability

The RNA-seq raw data generated in this study have been deposited at the accession XXX. The ChIP-seq raw data generated in this study have been deposited at the accession XXX. All the materials of the study are available from the Corresponding Author upon request.

## Acknowledgements

The authors thank Flavio Gabrieli for the helpful suggestions on bioinformatic analyses, and Enrico Magnani for kindly providing the *pABS:ABS-GFP* marker line and for useful discussion. We thank Marta Mendes, Cecilia Zumajo and Alex Cavalleri for the technical support and the stimulating discussion. Part of this work was carried out at the NOLIMITS platform, the advanced imaging facility of the Università degli Studi di Milano. This project was supported by Università degli Studi di Milano, the Humboldt-Universität zu Berlin, the University of Zurich, and the University of Adelaide. N.B. and C.A were supported by Università degli Studi di Milano Ph.D fellowship; N.B was supported by H2020-MSCA-RISE-2019 ID 872417; C.A. was supported by H2020-MSCA-RISE-2020; CB was supported by PRIN2017 and by PRIN2022.

## Author contribution

C.B., N.B., M.D.M., and L.C. conceived and designed the project. C.B., N.B., C.M., G.L. and M.D.M. carried out most of the experiments. R.V. performed and analysed the ChIP-seq experiment. F.A. performed the GS-MC metabolome profile. C.A. and M.D.M. analysed RNA-seq and ChIP-seq data. I.E., R.A.C, W.S., J.M.M., K.K., U.G., M.R.T., and L.C., supervised the work. C.B., N.B., M.D.M. and L.C. wrote the paper with assistance from all authors.

## Competing interests

The authors declare no competing interests.

## Supplementary Figures

**Supplementary Fig. 1.**
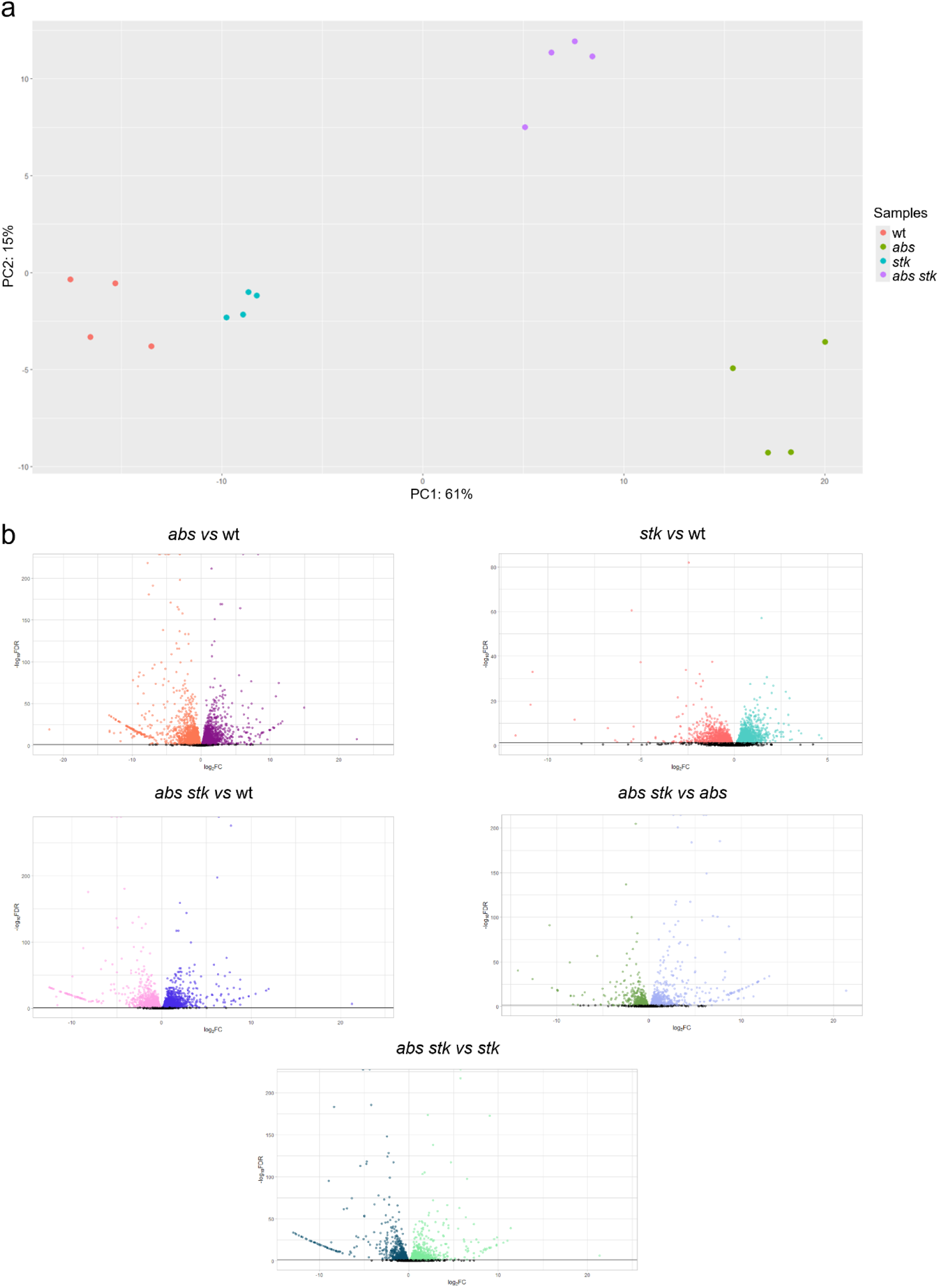
PCA and Volcano plot of RNA-seq data pre-fertilization. **a** Principal Component Analysis (PCA) displaying the first two principal components (PC1, PC2) shows good clustering of the replicates of the four different RNA-seq samples (wt, *abs*, *stk*, *abs stk*). **b** Volcano plots for the differential expression analysis showing distribution of DEGs in the different comparisons (*abs/*wt, orange: downregulated genes, purple: upregulated genes; *stk*/wt, red: downregulated genes, green: upregulated genes; *abs stk*/ wt, pink: downregulated genes, blue: upregulated genes; *abs stk*/*abs*, green: downregulated genes, light blue: upregulated genes; *abs stk*/*stk*, blue: downregulated genes, green: upregulated genes). n=4.

**Supplementary Fig. 2.**
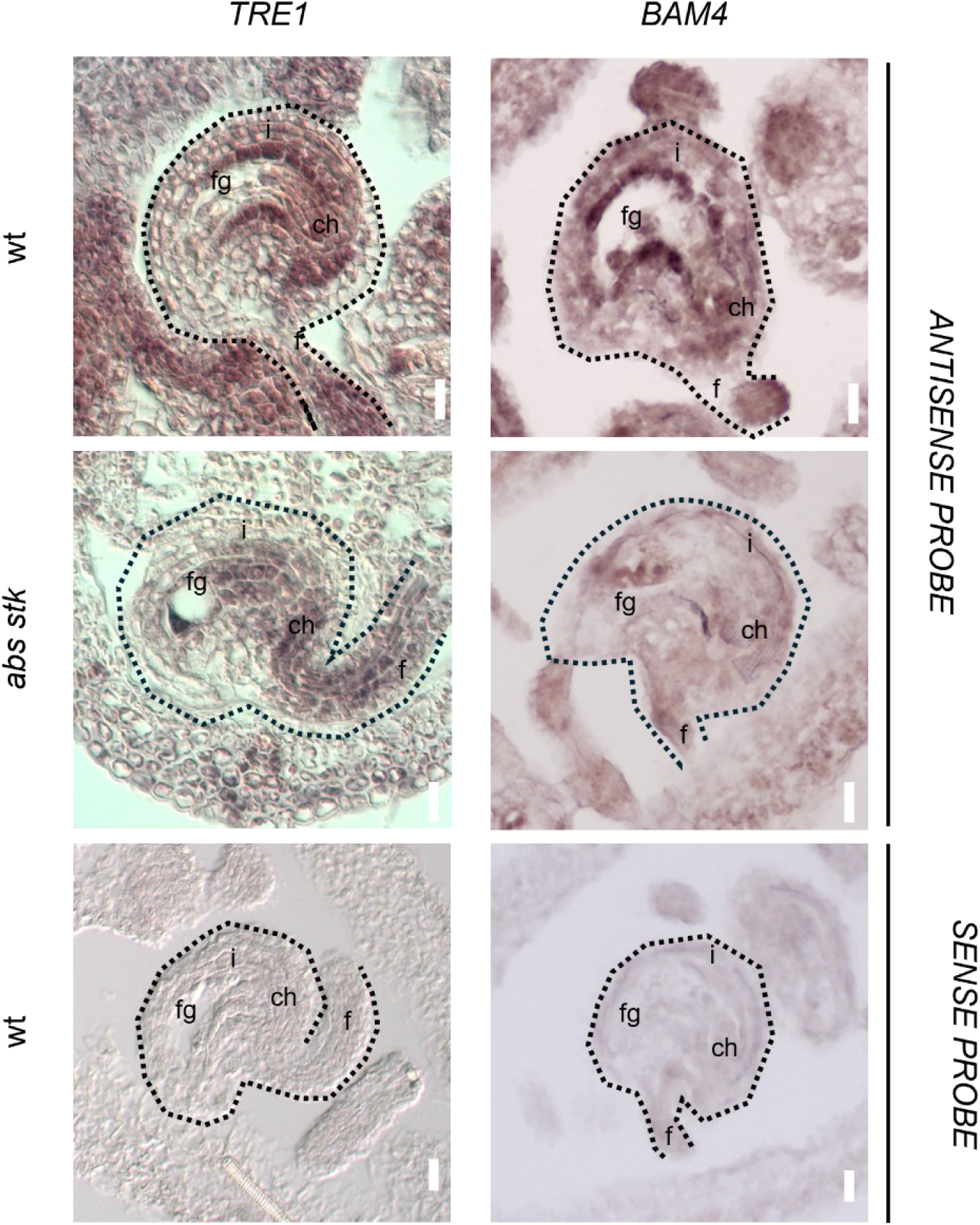
*In situ* hybridization on *TRE1* and *BAM4* in wt and *abs stk* ovules. The ovule border is highlighted by dotted line. fg: female gametophyte, f: funiculus, i: integuments, ch: chalaza. Scale bar 20 µm. Experiments were repeated on three independent individuals with similar results, and representative images are shown.

**Supplementary Fig. 3.**
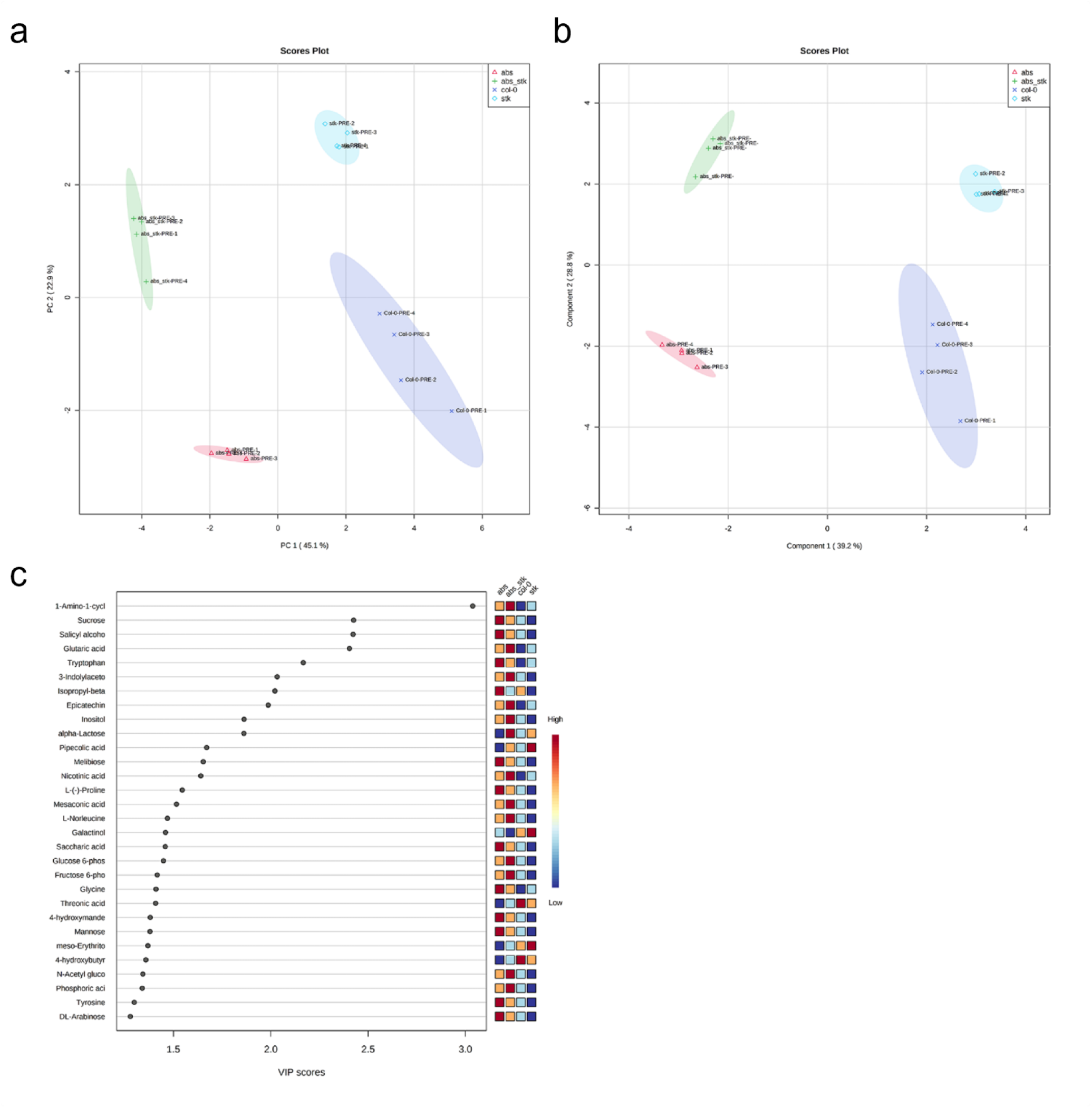
Different statistical analyses performed on the metabolome profile conducted on the wt and mutant flowers pre-fertilization. **a** PCA score plot, **b** PLSD-A score plot and **c** related VIP scores of wt (Col-0) and mutants (*stk, abs*, and *abs stk*) in Col-0 background. n=4.

**Supplementary Fig. 4.**
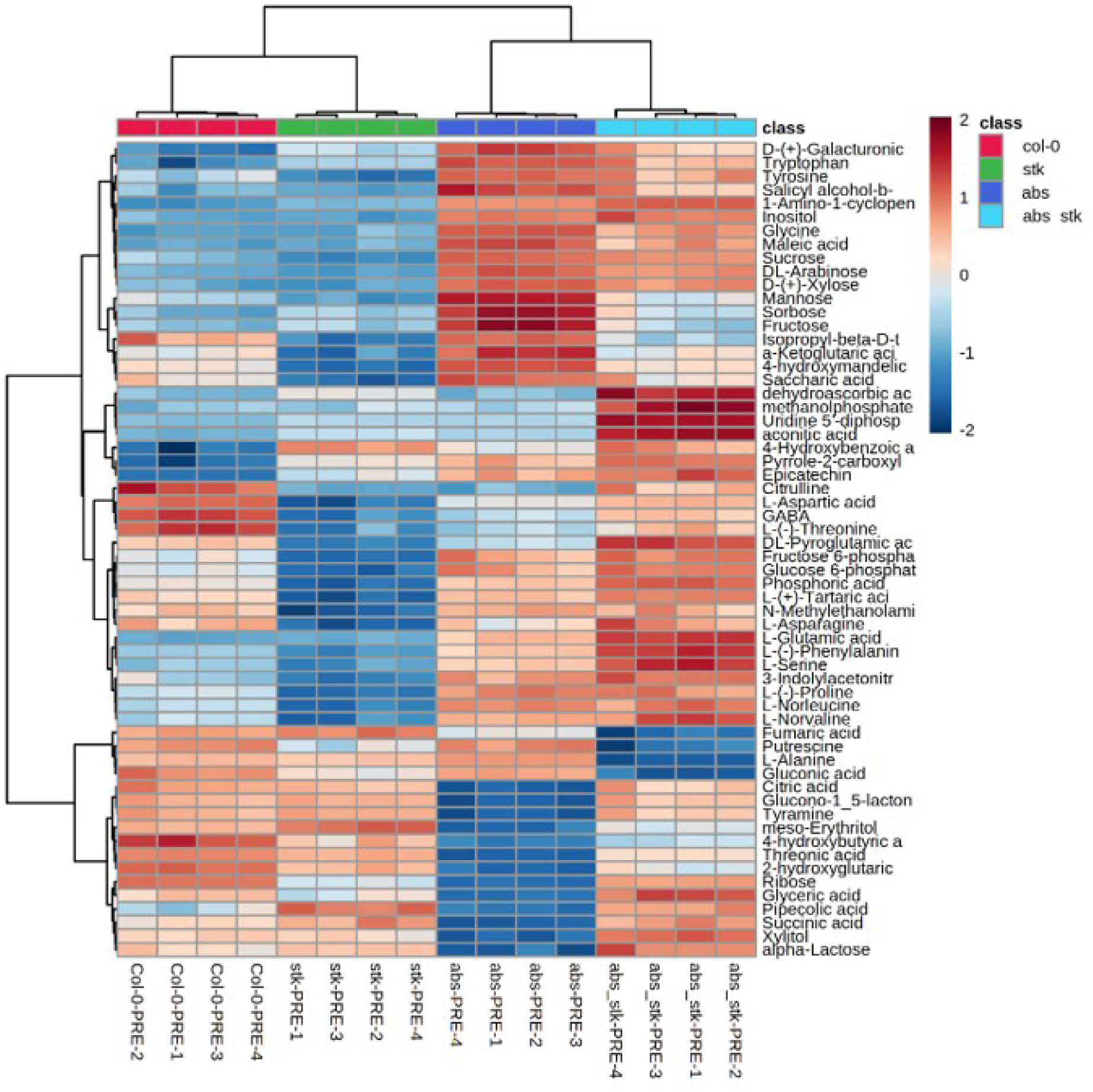
Heatmap showing differentially accumulated metabolites in wt and mutant plants. Clustering result shown as a heatmap built with the top 60 out of 122 metabolites with significant differences as determined by univariate one-way ANOVA using LSD test as post-hoc. Each rectangle represents the amount of a metabolite in false-color scale. Dark red or dark blue regions indicate an increase or decrease in metabolite content, respectively. Col-0: wt.

**Supplementary Fig. 5.**
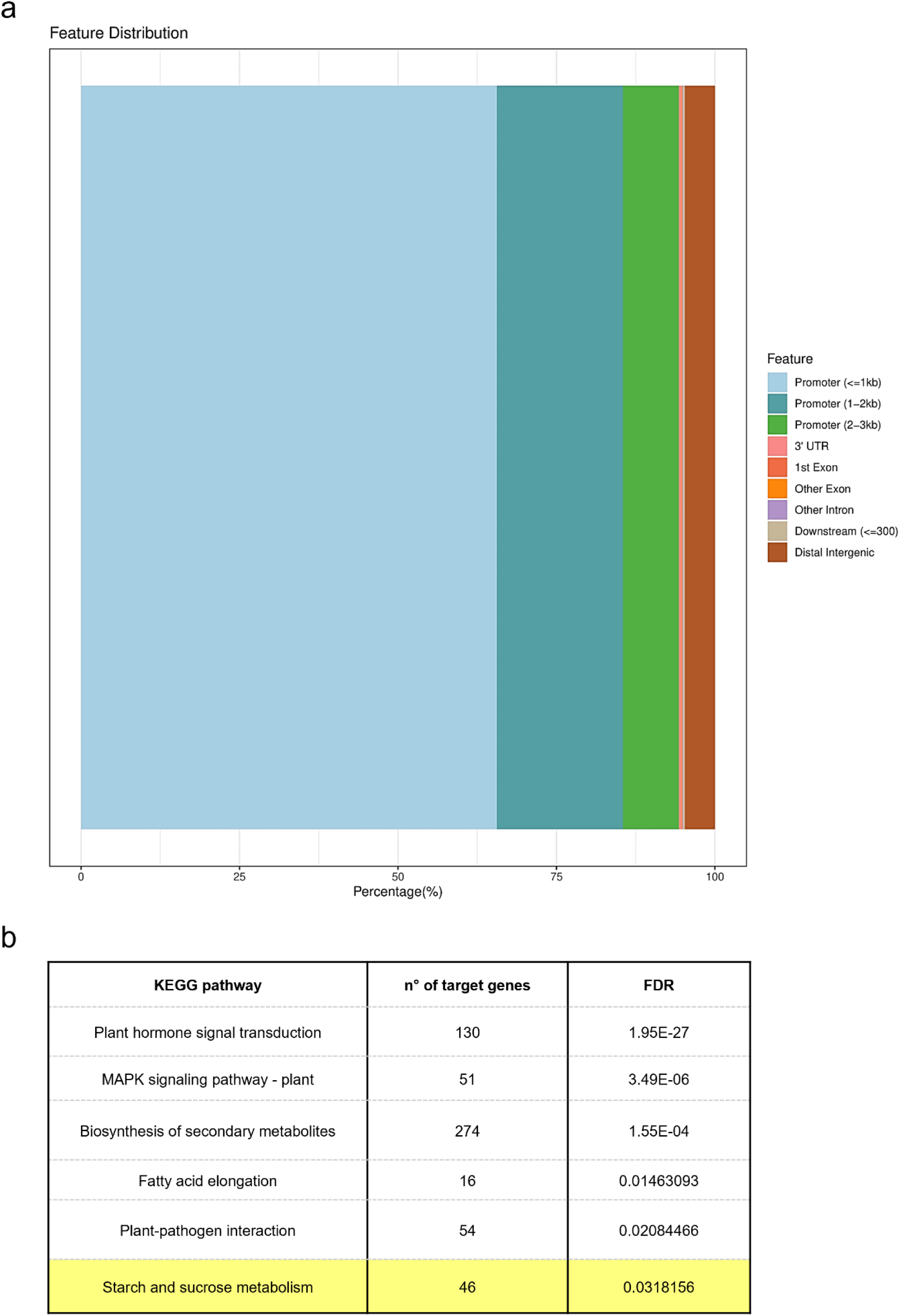
AnnoBar plot and KEGG enriched pathways in STK ChIP-Seq data pre-fertilization. **a** AnnoBar plot represents the percentage of peaks in the genomic region of target genes of STK. Most of the peaks are present in the promoter regions of STK target genes. **b** KEGG enriched pathways in STK ChIP-Seq data. Table representing the six most enriched KEGG pathways found on the STK ChIP-seq data with the respective number of STK target genes. Starch and sucrose metabolism category (highlighted in yellow) is enriched with 46 genes targeted by STK. Enriched pathways have been selected by an FDR < 0.05.

**Supplementary Fig. 6.**
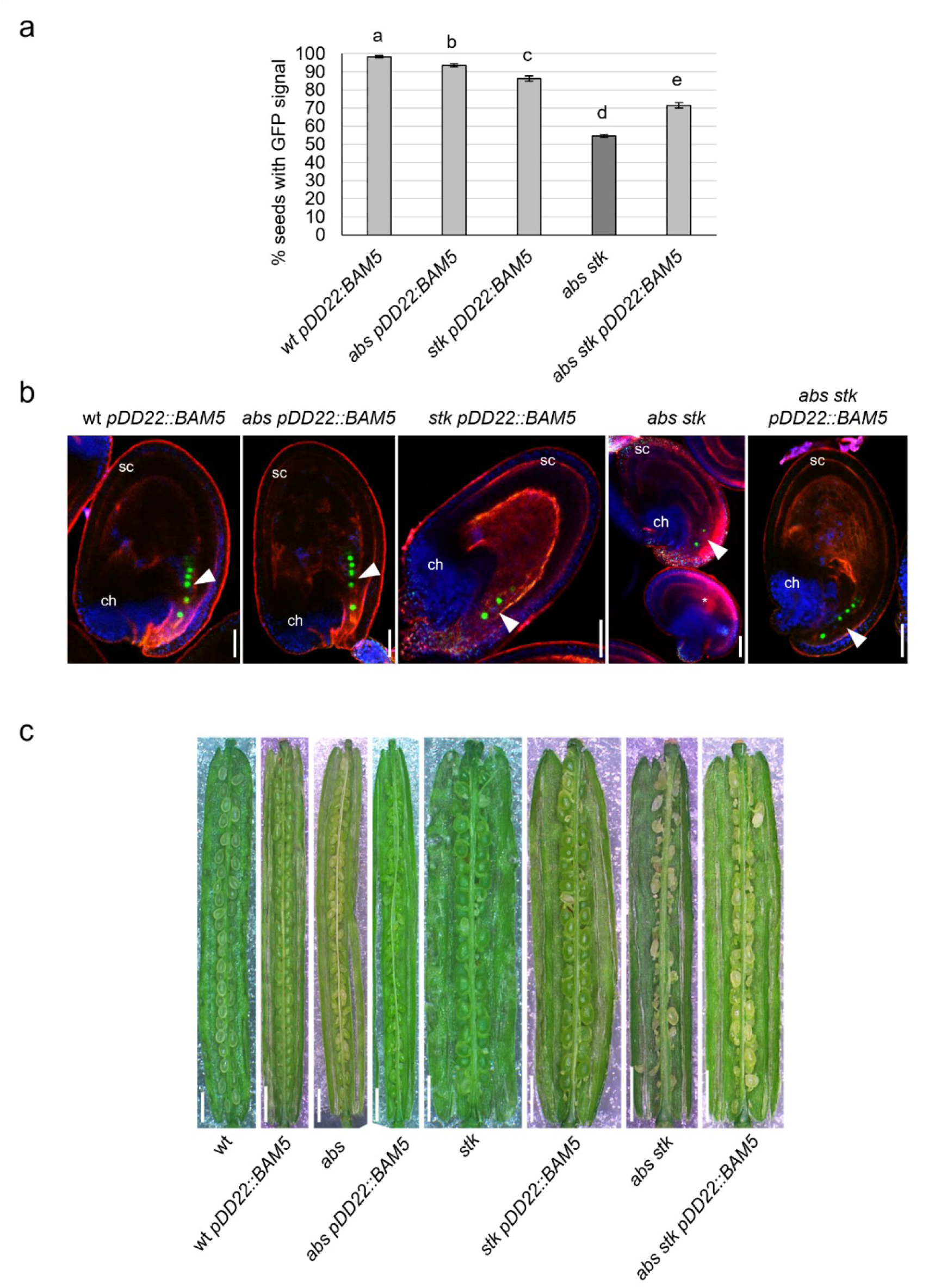
Characterization of the *pDD22::BAM5* transformant lines. **a** Percentage of seeds showing a WOX9-GFP signal at 3 DAP in wt *pDD22::BAM5* (n=13), *abs pDD22::BAM5* (n=13), *stk pDD22::BAM5* (n=11), *abs stk* (n=8), *abs stk pDD22::BAM5* (n=12). Error bars represent the standard error mean. Different letters above the bars represent significant differences (*p* < 0.05) as determined by one-way ANOVA followed by Tuckey’s HSD, whereas the same letters indicate absence of a statistical difference between the respective genotypes. Two independent measurements showed similar results and source data are provided as a Source Data file. **b** Confocal images of seeds of *pDD22::BAM5* transgenic lines expressing the WOX9-GFP protein in the suspensor of the embryos (arrowheads). **c** Siliques of the different transgenic lines and controls analyzed. ch: chalaza, sc: seed coat. Scale bars **b**: 50 µm **c**: 1mm. Experiments were repeated on at least three independent individuals with similar results, and representative images are shown in **b**, **c**.

**Supplementary Fig. 7.**
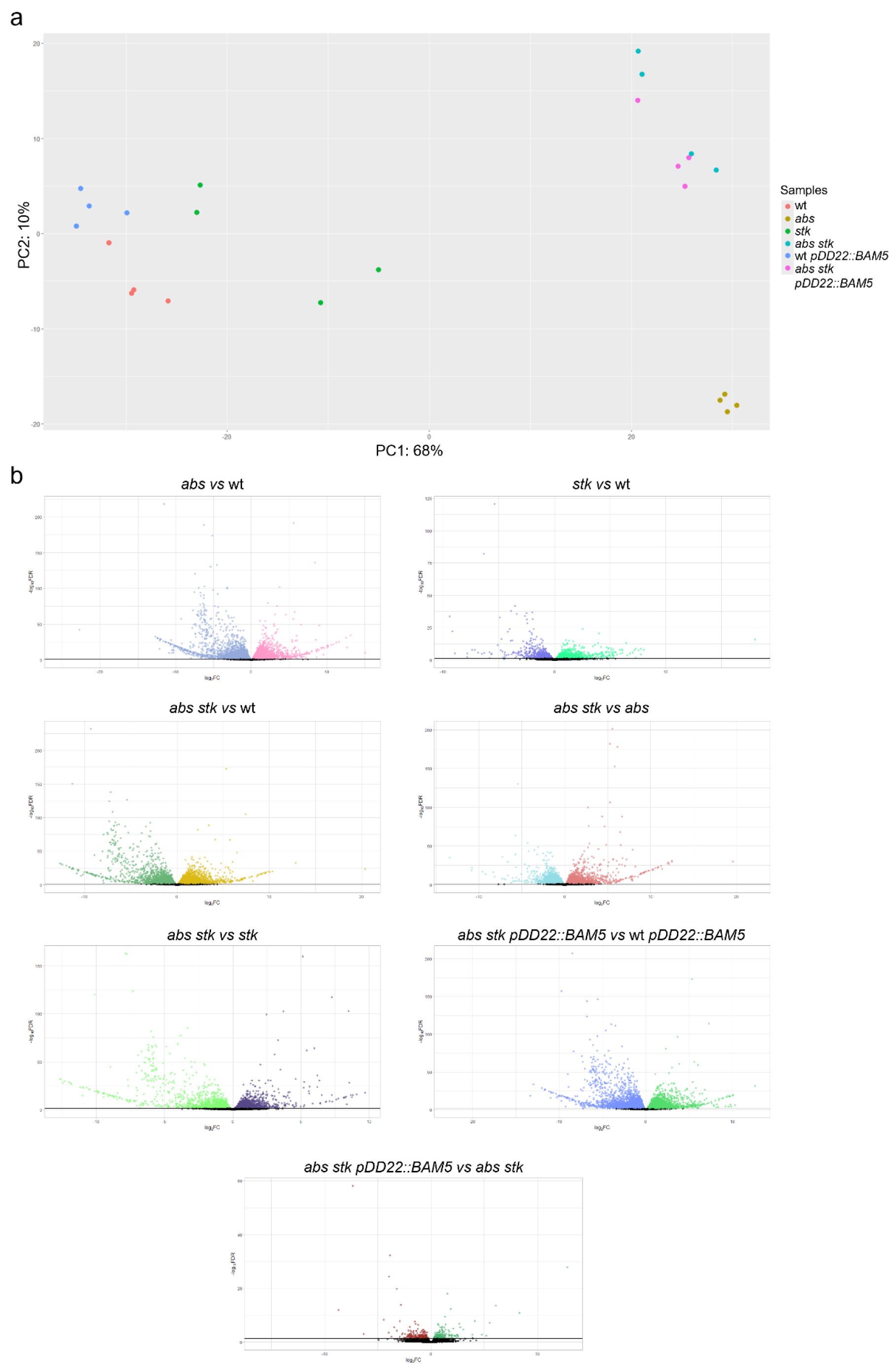
PCA and Volcano plot of RNA-seq data post-fertilization. **a** Principal Component Analysis (PCA) displaying the first two principal components (PC1, PC2) showing clustering of the four biological replicates of the RNA-seq samples at 3 DAP (*abs*, *stk*, abs *stk*, *abs stk pDD22::BAM5,* wt, wt *pDD22::BAM5*). n=4. **b** Volcano plot of the different comparisons for differential expression analysis shows distribution of DEGs (*abs/*wt, liliac: downregulated genes, pink: upregulated genes; *stk*/wt, blue: downregulated genes, green: upregulated genes; *abs stk*/ wt, green: downregulated genes, yellow: upregulated genes; *abs stk*/*abs*, light blue: downregulated genes, red: upregulated genes; *abs stk*/*stk*, green: downregulated genes, purple: upregulated genes; *abs stk pDD22::BAM5*/wt *pDD22::BAM5*, blue: downregulated genes, green: upregulated genes; *abs stk pDD22::BAM5*/*abs stk*, red: downregulated genes, green: upregulated genes).

**Supplementary Fig. 8:**
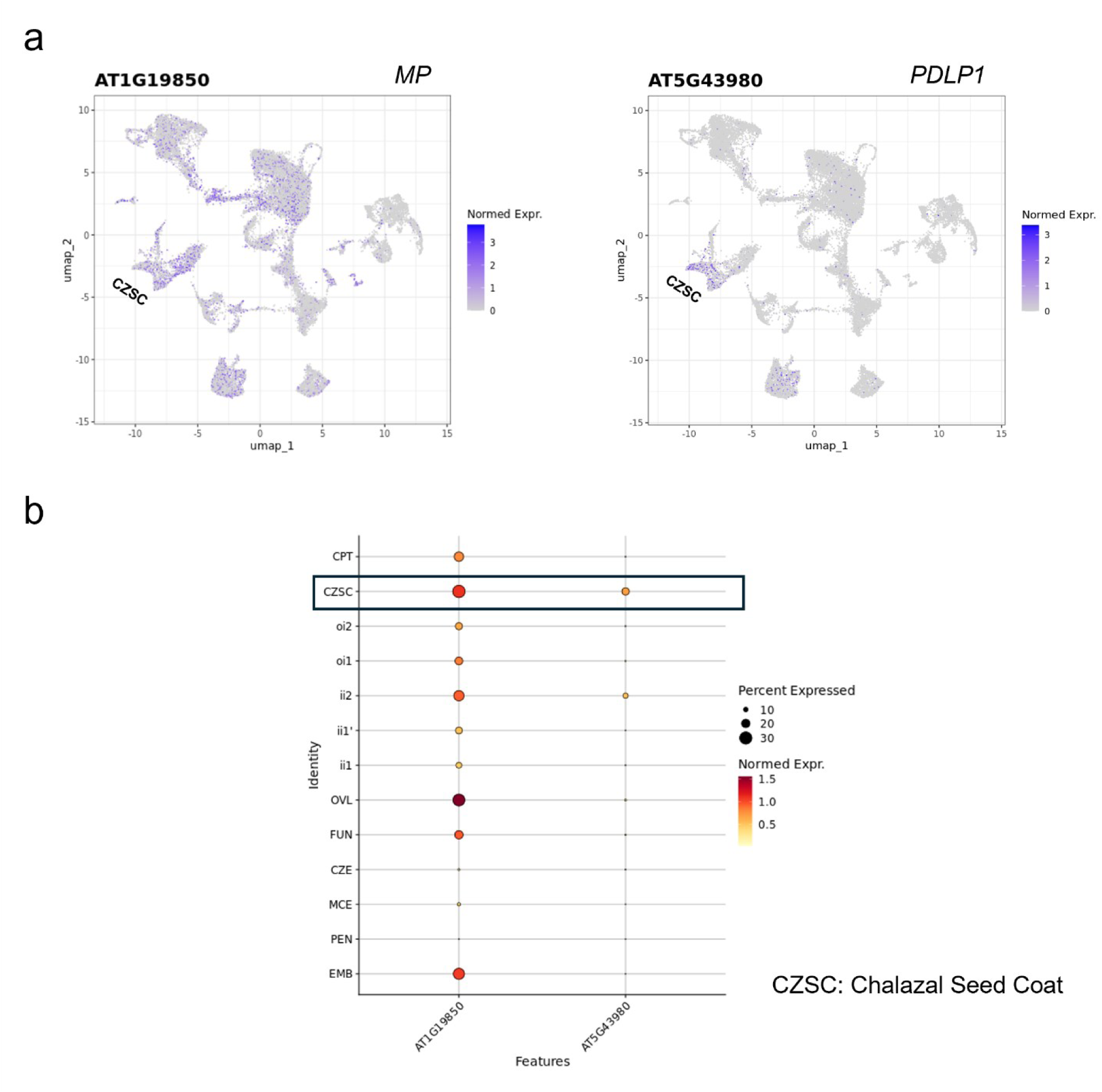
UMAP and Dotplot showing the expression pattern and level of *MP* and *PDLP1* in snRNA-seq from Martin et al., 2025. **a** Uniform Manifold Approximation and Projection (UMAP) showing the expression of AT1G19850 (*MP*) and AT5G43980 (*PDLP1*) in the seed at 3 DAP. **b** Dotplot showing the percentage of expression and the normalized expression of *MP* and *PDLP1*. Graphs are generated by an online database (Martin et al., 2025).

**Supplementary Fig. 9.**
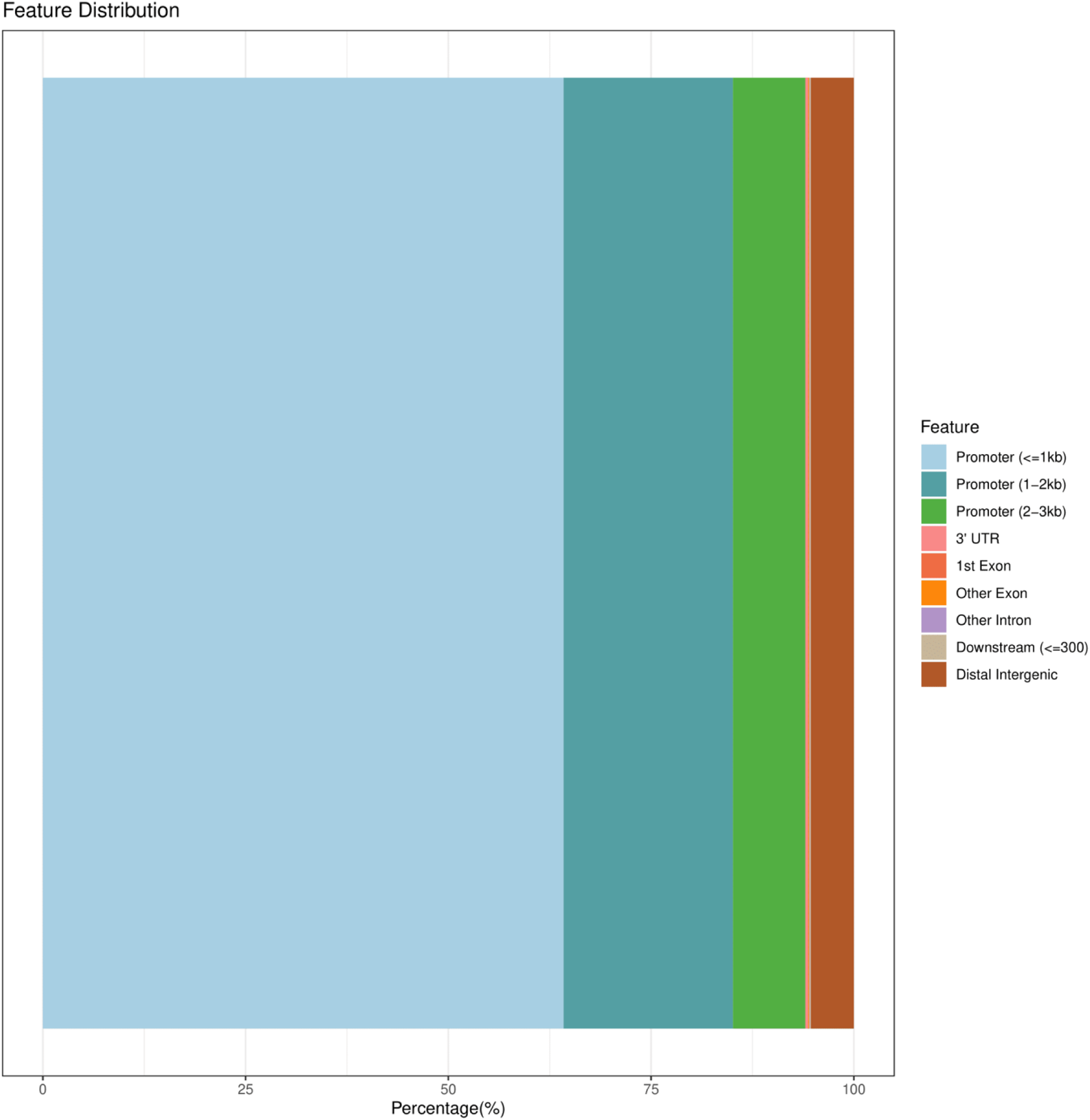
AnnoBar plot depicting the localization of the peaks in the STK ChIP-seq data post-fertilization. AnnoBar plot depicting the percentage distribution of ChIP-seq peaks across various genomic regions associated with STK target genes. Notably, a consistent proportion of the peaks are enriched in the promoter regions of STK target genes.

## Notes

### Competing Interest Statement

The authors have declared no competing interest.

